# Cerebrovascular pulsatility differs across vascular compartments and is altered by hypercapnic stimuli: a BOLD fMRI study

**DOI:** 10.64898/2026.06.09.730775

**Authors:** Hans Christian Rundfeldt, Wouter Schellekens, Emiel C.A. Roefs, Alex A. Bhogal, Mario Gilberto Báez-Yáñez, Jaco J.M. Zwanenburg, Natalia Petridou

## Abstract

Cerebral small vessel disease and neurodegenerative disorders have been associated with increased cerebrovascular pulsatility. Recently, BOLD fMRI-based methods have emerged for assessing pulsatility, however their interpretability is limited because the relation between estimated pulsatility indices (*PI*) and vascular anatomy and physiology remains poorly understood. To improve interpretability, we introduce a cardiac-specific BOLD fMRI– based *PI*, investigate its relationship to the cortical vasculature, and validate its sensitivity by introducing the known physiological vascular modulation of hypercapnia. Using high-resolution 7T BOLD fMRI with gradient-echo (GE) and spin-echo (SE) sequences, we disentangled macro- and microvascular contributions to the *PI* and quantified it across cortical depth. *PI* maps revealed anatomically plausible patterns, with elevated GE-*PI* near large veins and in white matter while SE-*PI* remained largely constant across cortical depth. GE-*PI* decreased during hypercapnia consistent with altered vascular tone, SE-*PI* on the other hand did not. *PI* correlated with cerebrovascular reactivity and venous blood volume suggesting sensitivity to vascular density and vessel mechanics. Our findings demonstrate that BOLD-derived *PI* provides a spatially and physiologically specific measure of vascular pulsatility. The BOLD fMRI-based *PI* method is readily applicable to existing datasets and has potential for assessing potential microvascular damage in cerebrovascular and neurodegenerative disease.

## 1. Introduction

Increased cerebrovascular pulsatility has previously been associated with the progression of vascular diseases in the brain such as small vessel disease (Delli Pizzi et al., 2023; Shi et al., 2018, 2020), and neurodegenerative disorders such as Alzheimer’s disease (Delli Pizzi et al., 2023; Roher et al., 2011). It is generally believed that larger arteries dissipate pulsatile energy, protecting fragile arterioles, capillaries, and venules from high pressure peaks (Zarrinkoob et al., 2016). Hereafter, we define the microvasculature as arterioles, capillaries, and venules embedded in the tissue. Arterial stiffening with aging or disease reduces the capacity of large vessels to buffer pressure pulsations. In turn, pulsatile energy propagates into the microvasculature potentially causing structural damage (Mitchell, 2008). Additionally, pulsatility has been proposed as a driving mechanism of glymphatic clearance (Iliff et al., 2013; Mestre et al., 2018; Rajna et al., 2021), suggesting that altered vascular pulsatility may impair waste removal from the brain.

Given its relevance as a potential disease mechanism, assessing changes of cerebral pulsatility may serve as an early biomarker for diverse pathologies. Pulsatility can be assessed brain-wide (e.g. by intracranial pressure monitoring [11]) or specifically for large blood vessels with respect to blood volume or blood flow velocity using phase-contrast MRI (PC-MRI) and transcranial doppler ultrasound (TCD) (Aaslid et al., 1982; van Hespen et al., 2022; Wagshul et al., 2011a). However, pathological changes predominantly affect the microvasculature (Pantoni, 2010), creating a spatial mismatch between the site of pathology and the vessels most accessible to measurements. This mismatch provides a challenge for diagnosing and understanding microvascular diseases.

To overcome this limitation and assess the microvasculature directly, refinements of existing techniques (e.g., PC-MRI (Arts et al., 2022; Pham et al., 2025; van den Kerkhof et al., 2023)) and novel approaches such as DENSE-MRI (Sloots et al., 2021; Van Hulst et al., 2024), vascular space occupancy (VASO) MRI (Guo et al., 2025), and velocity-selective arterial spin labeling (VS-ASL) (Chen et al., 2024) have been proposed. Several studies have employed blood oxygenation level–dependent (BOLD) fMRI to assess pulsatility as it offers high sensitivity to microvascular and venous signal changes (Atwi et al., 2020; Kim et al., 2021; Kiviniemi et al., 2016; Kornemann et al., 2023; Makedonov et al., 2013; Shirzadi et al., 2018; Sleight, 2023; Theyers et al., 2019; Tuovinen et al., 2020, 2024; Viessmann et al., 2017, 2019). However, the specificity of BOLD-derived pulsatility to the underlying signal-generating mechanisms and the underlying vascular anatomy, particularly with respect to vessel size, vascular density, and changes in vessel mechanics such as stiffening, remain poorly understood.

Therefore, this study investigates the dependence of the BOLD-based pulsatility index (*PI*) on vessel size, vascular density, and physiological changes in the vasculature. BOLD fMRI data were acquired at 7T using gradient-echo (GE) and spin-echo (SE) acquisitions with high spatial and temporal resolution. Given the microvascular specificity of SE BOLD versus the specificity of GE BOLD to both macro- and microvessels, we could compare GE-BOLD and SE-BOLD based *PIs* to characterize the *PI* dependence on vessel size (Budde et al., 2014; Duong et al., 2003; Uludağ et al., 2009). The high spatial resolution further allowed to assess *PIs* at different cortical depths to quantify pulsatility associated with different vascular densities and macro-/micro-vascular signal contributions. Finally, physiological changes in the vasculature were introduced via controlled hypercapnia and hyperoxia. Hypercapnia induces vasodilation and is hypothesized to alter vessel stiffness (Casaccia et al., 2014), serving as a model of altered vascular physiology in aging and disease and allowing to assess the sensitivity of the proposed pulsatility marker to changes in vessel-wall stiffening. Further, the hypercapnic challenge was utilized to estimate cerebrovascular reactivity (CVR), which describes the vasodilatory capacity of the vasculature in response to hypercapnia (Bhogal et al., 2016). Hyperoxia, by contrast, does not alter vascular tone (Chiarelli et al., 2007). However, hyperoxia increases venous oxygenation and thereby the BOLD signal. The resulting signal increase is utilized as a relative measure of (venous) cerebral blood volume (CBV). The relation of PI to CVR and CBV provided additional insights into the dependence of PI on vascular density and physiology.

## 2. Material and Methods

### 2.1 Subjects

We used data previously acquired at 7T as reported in (Schellekens et al., 2023). Eleven healthy individuals participated in this study after providing written informed consent. All participants reported no history of breathing difficulties under normal conditions and no known (cerebro)vascular-related illnesses. The study protocol was approved by the ethics committee at the University Medical Center Utrecht (UMCU), adhering to the principles of the Declaration of Helsinki (2013).

### 2.2 MRI Data

The 7T imaging protocol has been described in detail elsewhere (Schellekens et al., 2023). Briefly, anatomical images included a T1-weighted (T1w) volume acquired using MPRAGE with voxel size of 0.8×0.8×0.8mm^3^, TR/TE=7.0/2.97ms, field of view (FOV) of 40×159x159mm^3^ (anterior-posterior × inferior-superior × right-left), covering the visual cortex, and a proton density (PD) weighted volume with identical coverage. Additionally, three T2*-weighted (T2*w), flow-compensated anatomical volumes were acquired (3D-EPI, FOV 40×161×161mm^3^, voxel size of 0.5×0.5×0.5mm^3^, TR/TE=56/30ms). For this scan, magnitude and phase volumes were reconstructed.

Functional imaging of the early visual cortex was conducted with GE and SE echo-planar imaging (EPI) sequences. GE volumes were acquired with SENSE-factor=4.0, EPI-factor=31, water-fat chemical shift=30pixels, TR/TE=850/27ms, flip angle (FA)=50°, voxel size=1.0×1.0×1.0mm^3^, and FOV 7×128×128mm^3^. SE volumes were acquired with SENSE-factor=2.0, EPI-factor=63, water-fat chemical shift=45pixels, TR/TE=850/50ms, FA=90°, voxel size=1.5×1.5×1.5mm^3^, and FOV 7.5×190×190mm^3^. A larger voxel size was used for the SE scans to enhance the signal-to-noise ratio (SNR) (Panchuelo et al., 2015).

Per session, up to four fMRI time series were acquired (2xGE, 2xSE) with varying breathing protocols (see below). Each time series comprised 820 volumes, lasting 697s. Additionally, multiple volumes with reverse phase encoding direction were acquired for each GE and SE time series to correct for geometric distortions. Respiration was measured using a respiratory belt and the cardiac signal was monitored using a finger plethysmograph (PPU).

### 2.3 Physiological Data Processing

To determine the usability of an acquired PPU signal, the quality was assessed using two metrics: inter-beat interval (IBI) outlier fraction and the relative area under the curve (AUC) of the cardiac frequency band.

For the AUC metric, the power spectral density was estimated and the highest peak between 0.5Hz and 3.0Hz was assumed as the cardiac frequency *f*_*cardiac*_. The relative cardiac AUC was computed as the AUC around *f*_*cardiac*_ with a bandwidth of +/-0.2Hz divided by the total AUC between 0.1-5.0Hz. The IBI was computed as the time between two systolic peaks. If more than 30% relative cardiac AUC were found previously, mean IBI was computed as *IBI* = 1/*f*_*cardiac*_ and IBI limits were set to 50% and 150% of mean IBI, respectively. For lower relative cardiac AUC, the cardiac frequency was not determined reliably and absolute IBI limits of 0.5s-1.5s were enforced. Cardiac cycles were regarded to be IBI outliers if the IBI was outside of the prescribed limits. Subjects with a relative cardiac power of less than 30% or an IBI outlier fraction higher than 10% were excluded from further analysis (compare to supplement S15).

From the acquired respiration data, respiration volumes per time (RVT) were extracted (Birn et al., 2008).

### 2.4 Breathing Protocol

During functional imaging, participants were exposed to different gas-breathing conditions. A computer-controlled gas blender and sequential gas delivery system (3rd generation RespirAct™, Thornhill Research Inc., Toronto, Canada (Slessarev et al., 2007)) was used to modulate the concentrations of CO_2_ and O_2_ by targeting partial pressure end-tidal (Pet)CO_2_ and PetO_2_ levels. Baseline PetCO_2_ levels were measured individually for each participant before scanning. PetCO_2_ and PetO_2_ traces were recorded throughout the scan.

During each fMRI time series, four breathing stages were induced (Figure 1A):

**Figure 1:**
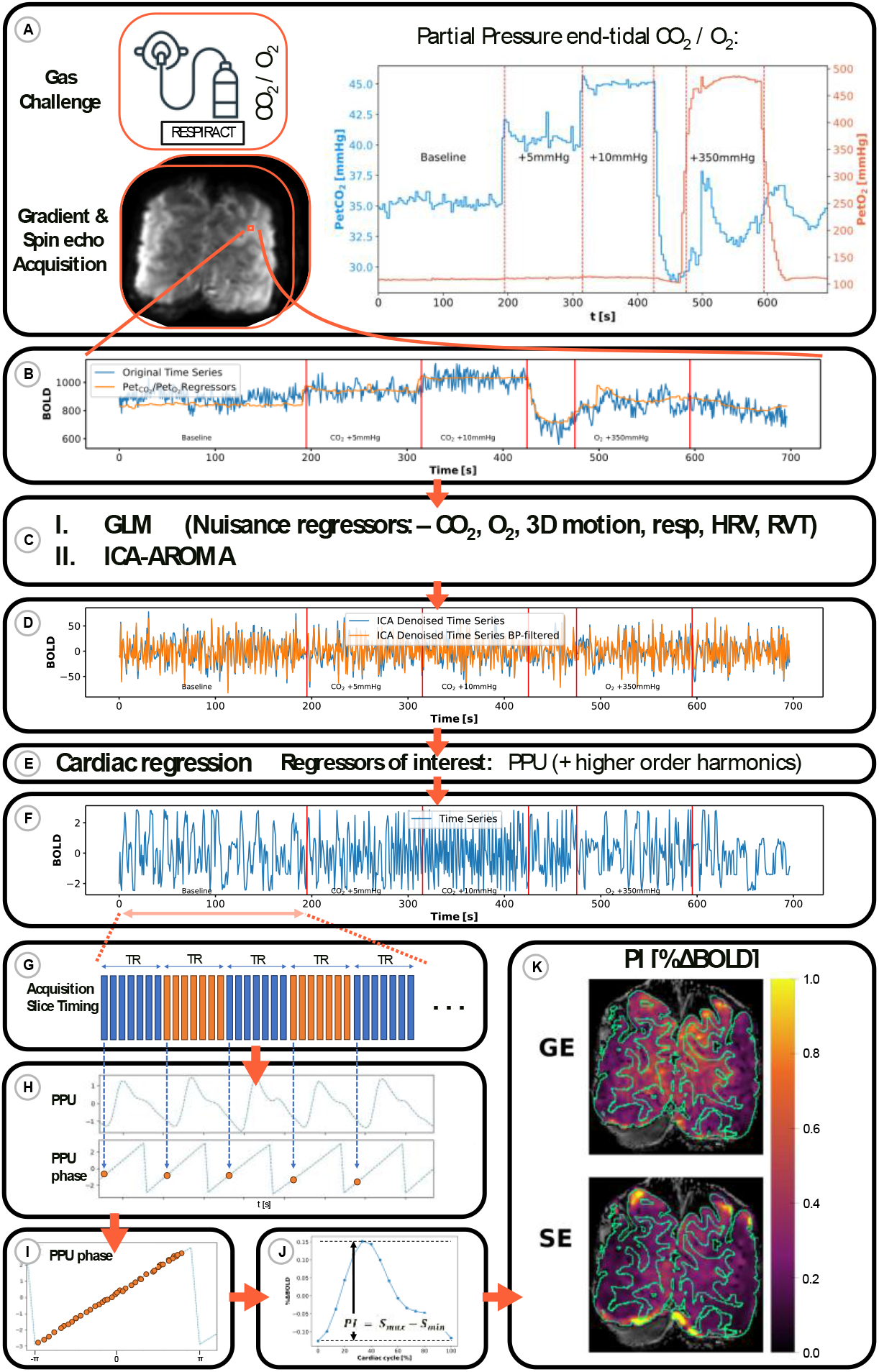
Pulsatility Index (PI) computation. A) Gradient-echo (GE) and Spin-echo (SE) BOLD fMRI scans were recorded separately, while the same breathing sequence was conducted as exemplified here. B) Example GE time series of one voxel (blue) and PetCO_2_/PetO_2_ regressors (orange). C) All nuisance regressors (motion parameters, respiration, HRV, RVT, and gas levels (PetCO_2_, PetO_2_)) were regressed out of voxel time series using a GLM. The residual voxel time series were further denoised using ICA-AROMA (Pruim et al., 2015). D) The denoised voxel time series (blue) were bandpass filtered to exclude low frequency oscillations (<0.1Hz, orange). E) A GLM with cardiac regressors from the PPU signal including the cardiac frequency and higher order harmonics was used to isolate cardiac components from the time-series. F) The resulting time series of isolated cardiac components represents signal changes due to cardiac pulsations. These cardiac time-series was separated into individual time series per gas level and normalized to percent signal change. G) BOLD fMRI acquired multiple slices per repetition (TR, alternating blocks are displayed in blue / orange). An individual voxels’ time series originates from one slice. H) The slice-timing aligned with the PPU acquisition was used to assign each time point a cardiac phase. I) The individual time series per gas level were resampled to one cardiac cycle based on the phase information. J) The resampled data points were averaged for 16 cardiac bins, yielding one signal over the cardiac cycle per voxel and gas level. PI is defined as the amplitude of the normalized signal per voxel. K) The presented workflow, repeated for GE and SE, yielded PI maps (here at baseline).

1. 200 s baseline: participant-specific PetCO_2_ targets.
2. 120 s hypercapnia: +5 mmHg PetCO_2_ increase at normoxia.
3. 120 s hypercapnia: +10 mmHg PetCO_2_ increase at normoxia.
4. 120 s hyperoxia: +350 mmHg PetO_2_ increase (at patient-specific normocapnic PetCO_2_ targets).

This breathing task was repeated, for GE and SE acquisition, respectively. Throughout the acquisitions, visual stimuli were presented at random times. Results related to the visual stimuli were previously reported and were excluded in this work by treating the respective hemodynamic response functions (HRF) as nuisance regressors (Roefs et al., 2024).

### 2.5 Structural Data Preprocessing

The T1w volume was normalized with the PD-weighted volume to correct for intensity inhomogeneities. The three T2*w volumes were realigned and averaged. Veins were segmented from T2*w magnitude images using Braincharter (Bernier et al., 2018) and quantitative susceptibility images derived from the T2*w phase images using Nighres (Huntenburg et al., 2018), and the union of both estimated masks was taken. The final vein masks were transformed from T2*w space to the respective functional space and resampled to probability masks (AFNI 3dfractionize (research & 1996, 1996)). The process of vein segmentation was previously described in detail [40].

T1w and T2*w scans were aligned using an affine transformation (AFNI 3dAllineate (Saad et al., 2008)), and the T1w scan was upsampled to the T2*w scan’s resolution using cubic interpolation. Using both aligned volumes, a two-channel tissue segmentation was conducted using SPM12 (Penny et al., 2007). The resulting tissue probability maps were transformed and resampled to the respective resolution of the GE and SE volumes using linear interpolation, and binarized to gray matter (GM) and white matter (WM) masks including voxels with a probability greater than 0.7. Twenty cortical GM laminae were estimated from the upsampled T1w volume (see (Schellekens et al., 2023)) using LayNii (Huber et al., 2021). The 20 individual laminae were summarized to three distinct laminae of deep, middle, and top GM. Any voxels with a vein probability greater than 0.1 were excluded from the laminae and the tissue masks. Resulting regions of interest (ROIs) are presented in Figure 2. Probability masks of the laminae were computed when registering and downsampling them to functional space to reduce the partial-volume effects (AFNI, 3dfractionize).

**Figure 2:**
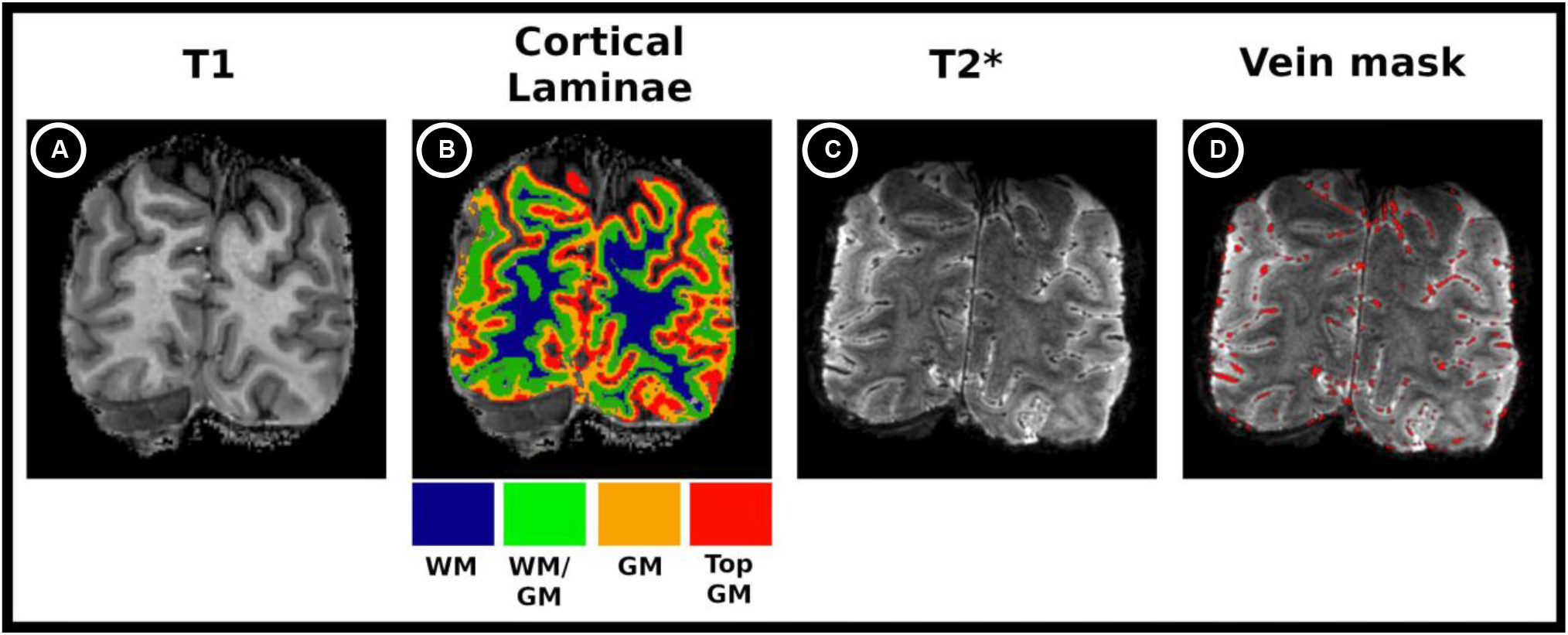
Structural Data Processing. A) T1w anatomical image. B) Cortical laminae overlayed to the T1w image with white matter (blue), deep gray matter (green), middle gray matter (orange), and top gray matter (red). C) T2*w anatomical image. D) Cortical vein mask overlayed to the T2*w image.

### 2.6 Functional Data preprocessing

Functional data were in-plane motion-corrected (AFNI 2dImReg), distortion-corrected using opposing phase-encoding volumes (AFNI 3dQwarp), and registered to T2*w anatomy (AFNI/ANTs, best result chosen). Anatomical masks were then transformed into functional space using the inverse transforms, leaving functional data in its native space. Cerebrovascular reactivity (*CVR*^*GE*/*SE*^) and oxygen-concentration based venous CBV 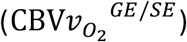 were estimated from hypercapnia and hyperoxia challenges, respectively, where the subscript 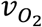 reflects the signal’s dependence on venous oxygenation and the superscript the scan sequence used. The procedure was previously reported (Roefs et al., 2024). *CVR*^*GE*/*SE*^and 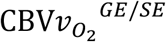 are reported as the regression coefficients between the change in BOLD signal (%ΔBOLD) and the PetCO_2_ / PetO_2_ gas trace and averaged per ROI. GLM t-statistics for hypercapnia and hyperoxia were corrected for false discovery rate (FDR, threshold q < 0.05) and responsive voxels were selected with a t-statistics threshold (t > 1).

### 2.7 Pulsatility Index Computation

The pulsatility index (*PI*) was computed per voxel. The procedure is illustrated in Figure 1. A GLM was applied to exclude nuisance regressors (3D motion parameters (AFNI 3dvolreg), respiratory belt data, visual task HRF, RVT, PetCO_2_/PetO_2_-traces) from the BOLD fMRI data Figure 1C). The GLM analysis was conducted per imaging slice with regressors offset by the slice timing. Residual time-series were further denoised using ICA-AROMA, which decomposes data into independent components and excludes those that correlate temporally with estimated motion parameters and spatially occur in voxels proximal to sudden transitions in image intensity (Pruim et al., 2015). The denoised time-series were high-pass filtered (>0.1Hz) to exclude low frequency fluctuations and signal drift (Balduzzi et al., 2008) (Figure 1D). Cardiac contributions were separated from the resulting time-series using RETROICOR as described by Glover et al. (Glover et al., 2000) (Figure 1E). This approach models the cardiac and respiratory components using a 2^nd^ order Fourier series where the basis functions are defined by the cardiac and respiratory phases at each acquisition time point. The modelled cardiac signal is least-square fitted to each voxel’s time series to recover the aliased cardiac waveform, allowing to analyze cardiac contributions in undersampled data. Next, the time-series was split into epochs for the baseline period, the two breathing stages of hypercapnia (+5mmHg and +10mmHg PetCO_2_), and the hyperoxia stage (+350mmHg PetO_2_, Figure 1F). These separate time-series were normalized, and retrospectively gated into 16 cardiac bins (Figure 1F-J).

Let *S*_*cardiac*_(*t*) and *S*′_*cardiac*_(*t*) be the cardiac time series before and after the retrospective gating. Further, 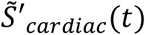 is the normalized gated cardiac time series, where tilde indicates the normalization (subtracting and dividing by its temporal mean 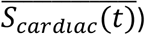. The peak-to-peak range of 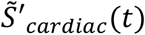 per voxel is defined as the pulsatility index according to 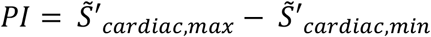, where 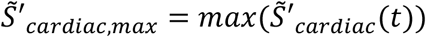 and 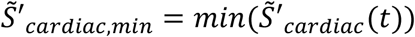. Similarly, the pulse amplitude (*PA*) per voxel is *PA* = *S*′_*cardiac,max*_ − *S*′_*cardiac,min*_ . Of note, the *PI* can be computed equivalently by 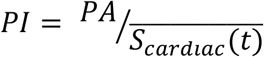. Exemplary resulting *PI* maps for GE and SE are illustrated in Figure 1K. In the following, *PI* and *PA* assessed by GE or SE are indicated with a superscript (*PI*^*GE*/*SE*^ or *PA*^*GE*/*SE*^) when necessary.

Mean *PI*^*GE*/*SE*^ were computed per laminae ROI by computing the voxel-wise *PI*^*GE*/*SE*^ followed by the spatial ROI-based average. Additionally, the computation was repeated after temporal alignment of the cardiac gated voxel time-series. The alignment was performed by shifting all voxel time-series to a reference cortical GM voxel with high temporal SNR using the offset that resulted in the highest temporal correlation with the reference voxel. For comparison, a mean cardiac time-course per ROI was computed and the 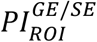 of the ROI was computed from the mean aligned cardiac time-course (see supplementary material S2). For visual comparison of *PI*^*GE*^ and *PI*^*SE*^, the resulting maps were transformed back to T2*w space with linear interpolation.

PetCO_2_ and PetO_2_ traces were decomposed into within-subject (ws) and between-subject (bs) changes. Within-subject changes allow to assess PI changes due to changed gas levels within a subject during a scan. To quantify this change, a voxel-wise linear regression between *PI*^*GE*/*SE*^ and within-subject hypercapnia / hyperoxia levels was conducted. The resulting slopes, 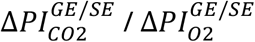 indicate the level of *PI*^*GE*/*SE*^ change within a subject following hypercapnia / hyperoxia stimuli. As the targeted end-tidal gas levels are not always exactly matched in the experiment, the mean gas levels per breathing condition are used rather than the target values. Between-subject changes account for different baseline signal levels correlating to different baseline end-tidal gas levels.

### 2.8 Statistical Analysis

Linear mixed-effects models (LMMs) were fitted to the data, allowing to include all voxels as individual observations. To equalize ROI sizes and assert that results are not dominated by individual subjects, the data was bootstrapped (n=10,000). For each subject and condition (scan sequence × breathing stage x laminae), a random sample of observations equal to the group mean number of ROI data points was drawn and averaged, yielding the ROI mean number of data points per subject, laminae, and scan type.

Model 1 assesses the effects of scan sequence and cortical depth (laminae) on *PI*. The laminae and scan sequence were chosen as categorical fixed-effect variables. Within- and between-subject gas levels were included to assess PI differences attributable to different gas levels. Interactions of scan sequence and within-subject gas levels were added to account for different responses to altered breathing gas mixtures for GE and SE, respectively. Additionally, an interaction of scan sequence and laminae was modelled to assess if *PI* behaved differently across laminae for GE and SE. Model 1 was fitted on all data points.

#### Model 1 (*PI* vs. scan sequence and laminae)

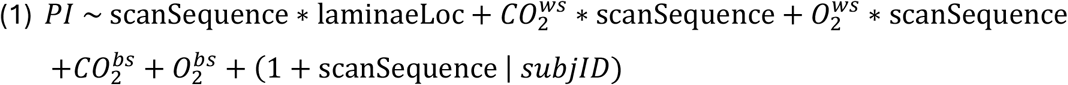

Model 2 assesses the spatial and scan sequence dependence of the PI change with hypercapnia (Δ*PI*_*CO*2_). Again, an interaction of scan sequence and laminae was modelled. The same test was performed for Δ*PI*_*O*2_ as a control analysis, as no change in pulsatility levels was expected in response to hyperoxia, and reported in the supplementary material (compare to supplement S14).

#### Model 2 (Δ*PI*_*CO*2_ vs. scan sequence, laminae)

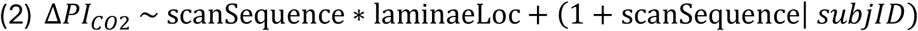

Further, the relationship of *PI* at baseline (Model 3, 4) and Δ*PI*_*CO2*_ (Model 5, 6) to *CVR*^*GE*/*SE*^ and 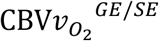 were modelled in separate LMMs for *CVR*^*GE*/*SE*^ and 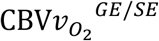, respectively. Voxels with 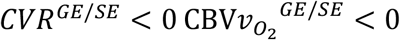 were excluded. The laminae and scan sequence and their interactions with *CVR*^*GE*/*SE*^ and 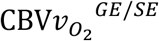 were included as additional fixed-effect variables. The interactions assess whether PI and Δ*PI*_*CO2*_ correlate with CVR differently for different scan sequences and laminae, respectively. Random slopes of *CVR*^*GE*/*SE*^ and 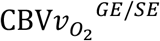 were estimated per participant.

#### Models 3-6 (*PI*/Δ*PI*_*CO*2_ vs. *CVR*/*CBV*)

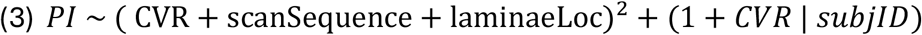

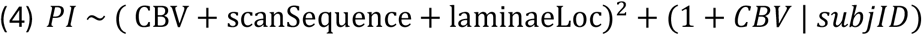

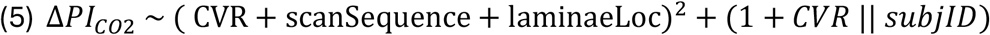

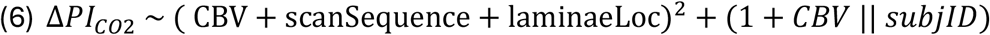

In all LMMs, a random intercept and slope per subject was fitted to account for varying baseline PIs in subjects. If singularity issues arose, the random structure was simplified by removing the intercept-slope correlation.

All models were fitted with restricted maximum likelihood (REML) using the *lmerTest* package (R v4.5.1) with optimizer *bobyqa* (Kuznetsova et al., 2017). Given the large number of observations but small number of subjects, Satterthwaite approximations of degrees of freedom were unreliable. Therefore, the models were fitted on aggregated data, and inference on fixed effects was performed using parametric bootstrap likelihood ratio tests (LRT) (*pbkrtest*, 10,000 simulations (Halekoh & Højsgaard, 2014)). The test statistic (LRT) and the bootstrap p-value (PBtest) are reported. Model fit visualizations are provided in S5, and full model specifications in S6-S13.

Post-hoc tests were performed on aggregated data using the *emmeans* R package, with degrees of freedom approximated using the Kenward–Roger method and multiple comparisons adjusted using the Tukey procedure.

## 3. Results

Of eleven participants, one was excluded due to incomplete scans and three due to low PPU quality (compare Supplement S15), leaving seven subjects (N=7; age 25–31 years; mean 29.1).

### 3.1 Pulsatility Index Maps

Figure 3 shows the *PI*^*GE*^ and *PI*^*SE*^ maps of a representative subject. The first column displays the baseline condition, followed by the hypercapnia levels (+5mmHg PetCO_2_ and +10mmHg PetCO_2_). *PI*^*GE*^ visually appears higher in WM vs GM across all gas levels, whereas only few high *PI*^*SE*^ voxels appeared in WM. Towards the brain surface and bordering the sagittal sinus, higher *PI*^*GE*/*SE*^ values were observed, with high *PI* voxels collocating between sequences. More local *PI*^*GE*^ peaks were visible, though *PI*^*SE*^ peaks expressed larger values.

**Figure 3:**
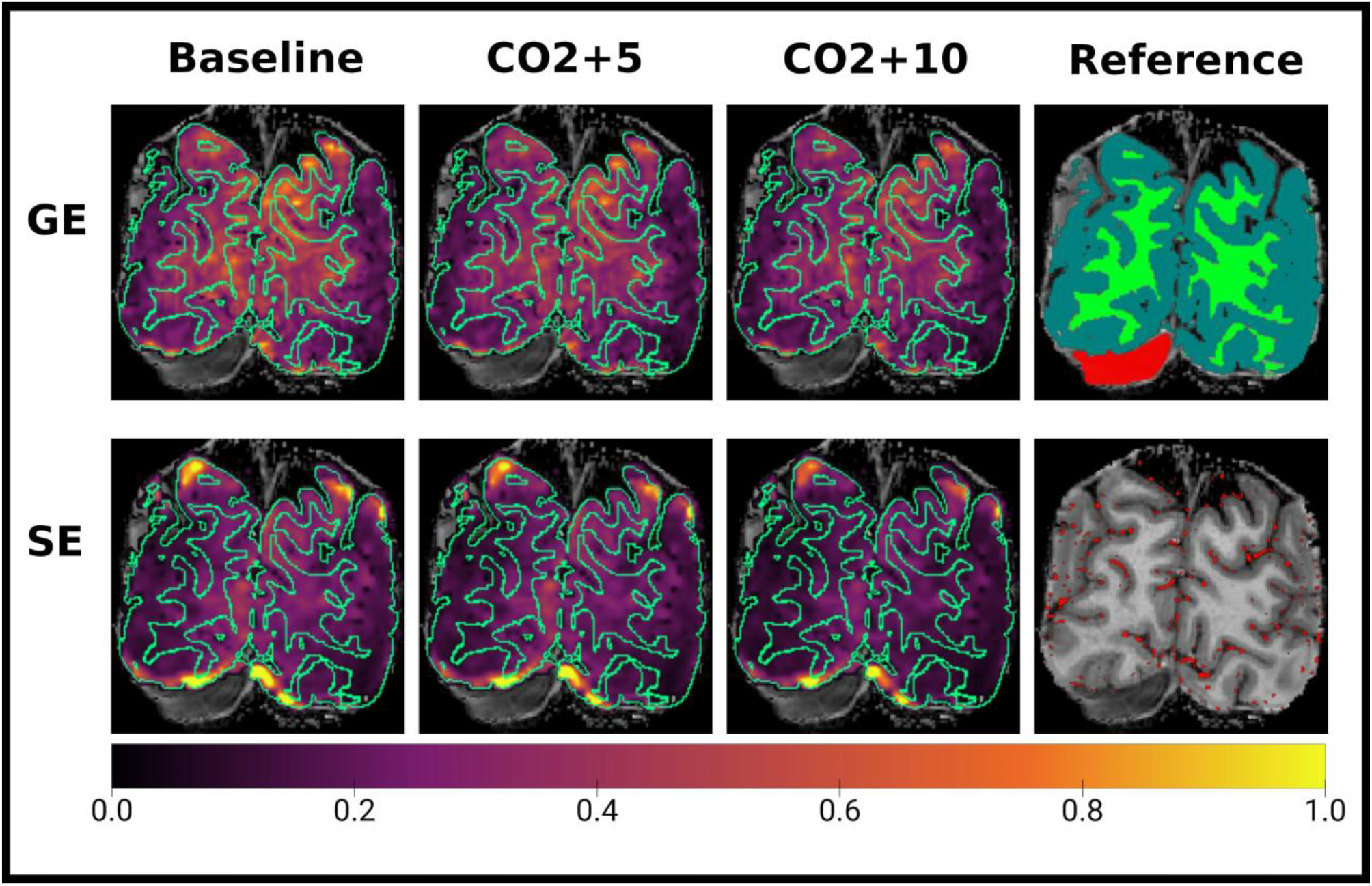
Pulsatility Index (PI) Maps. From left to right: PI estimated using Gradient-echo (Top row) and Spin-echo (bottom row) at baseline, +5mmHg PetCO_2_, and +10mmHg PetCO_2_, respectively. The right most column shows the gray matter segmentation (dark green), the white matter segmentation (light green), and the straight sinus (red) in the top row. In the bottom row, the vein mask is displayed.

With increasing hypercapnia levels, a decrease in *PI*^*GE*/*SE*^ is visible, which is stronger for *PI*^*GE*^ than for *PI*^*SE*^.

### 3.2 Pulsatility Index across Cortical Depth

Figure 4 displays ROI-averaged *PI*^*GE*/*SE*^, *PA*^*GE*/*SE*^, and mean cardiac signal per subject across gas levels.

**Figure 4:**
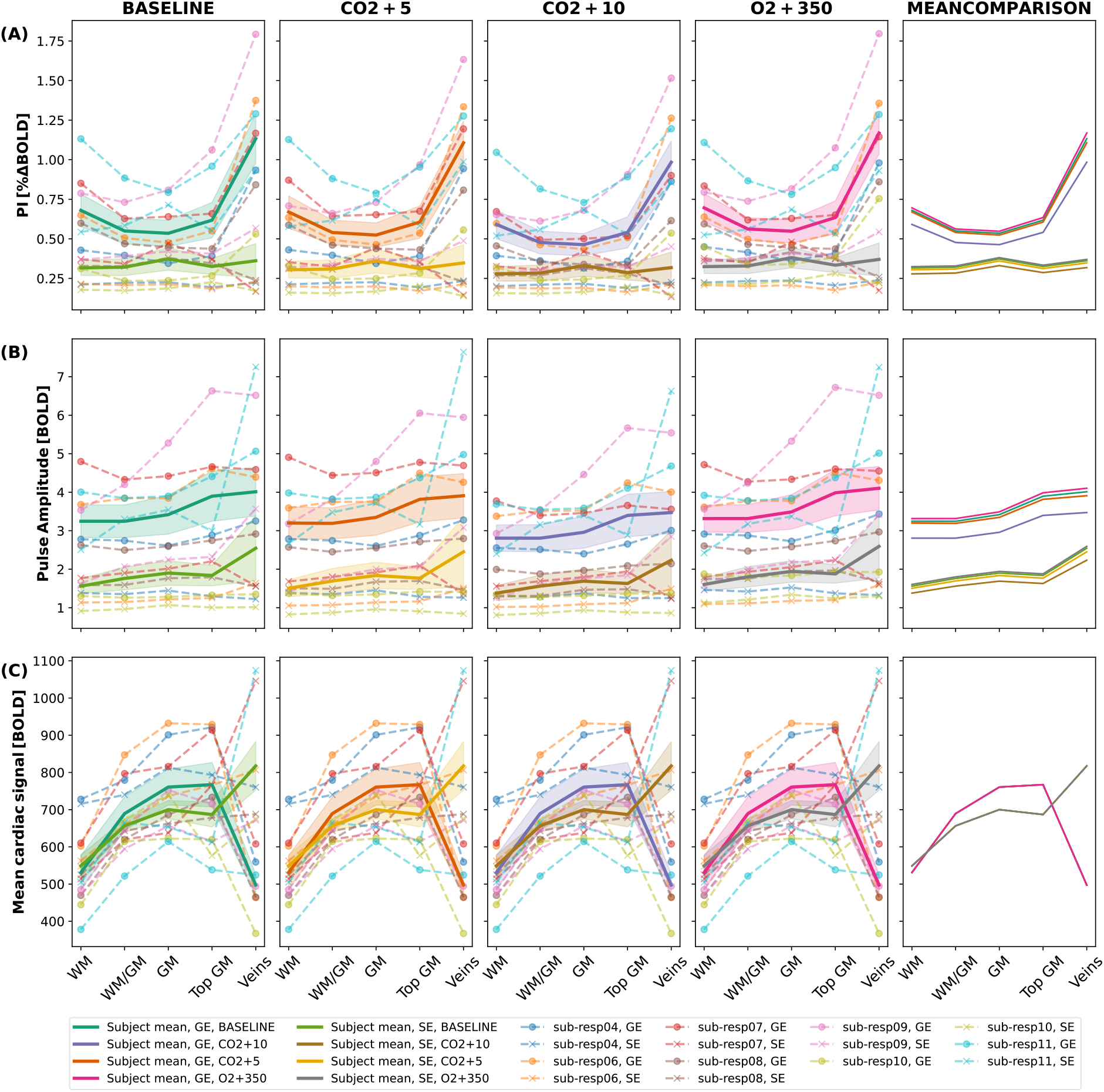
Pulsatility Index (PI [%ΔBOLD]), Pulse Amplitude (PA [BOLD]), and mean signal of the ungated cardiac-filtered time series (Mean cardiac signal [BOLD]) across cortical depth. The pulsatility index (top row, A), pulse amplitude (middle row, B), and mean cardiac signal (bottom row, C) are plotted across cortical depth at baseline, +5mmHg PetCO_2_, +10mmHg PetCO_2_, and +350mmHg PetO_2_ (left to right). Mean quantities per ROI are plotted for the individual subjects (dashed lines) and the subject average (thick lines, shaded area represents standard error of mean) for GE and SE. The subject averaged cortical depth profiles from all breathing conditions are plotted in the right most column for visual comparison (only one curve is visible in the for the Mean cardiac signal due to overlap). The PI (top row) is obtained by normalizing the PA (middle row) with the mean cardiac signal (bottom row).

During all breathing conditions, *PI*^*GE*^ and *PI*^*SE*^ differed significantly (Model 1: 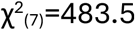, p=.0001). Cortical depth significantly explained variance (Model 1: 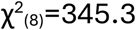, p=.0001) and a significant scan sequence and cortical depth interaction indicated differing spatial distribution of *PI*^*GE*^ and *PI*^*SE*^ between sequences (Model 1: 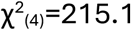, p=.0001). *PI*^*GE*^ were lowest in the middle GM and increased towards the pial surface (post-hoc, t_(256)_=-22.2, p<.0001) and the WM (post-hoc, t_(256)_=5.4, p<.0001). Contrary, *PI*^*SE*^ expresses a visible peak in the middle GM, however no significant decrease towards WM (post-hoc, t_(256)_=-2.1, p=0.22) and top GM (post-hoc, t_(256)_=0.47, p=0.99). The linear mixed model coefficients, PBtest values, and post-hoc tests can be found in supplement S6.

*PA* distributions (Figure 4, second row, left column) differed from *PI* . *PA*^*GE*^ was less increased in large veins compared to top GM and did not increase in WM relative to middle GM. *PI*^*GE*^ and *PA*^*GE*^ differences reflected the normalization by the mean GE cardiac-BOLD signal (Figure 4, third row) which was low in WM and large veins compared to GM laminae. Conversely, the mean SE cardiac-BOLD signal peaked in large veins and was lowest in the WM, yielding higher *PA*^*SE*^ in large veins and lower *PA*^*SE*^ in WM compared to *PI*^*SE*^ . The full linear mixed model coefficients and PBtest values for the *PA* and the mean cardiac signal can be found in the supplementary material (S7, S8). The relationship between voxel PA and mean signal is plotted in the supplementary material (S3).

### 3.3 Pulsatility Index Change during Hypercapnia and Hyperoxia

The four columns of Figure 4 show ROI-averaged *PI*^*GE*/*SE*^ at baseline compared to hypercapnia (PetCO_2_ +5mmHg, +10mmHg) and hyperoxia (PetO_2_ +350mmHg). Across all laminae, *PI*^*GE*/*SE*^ levels varied significantly with PetCO_2_ within-subject (Model 1: 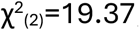, p<.0001), but not between-subject (Model 1: 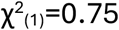, p=0.6). *PI*^*GE*^ decreased at +10mmHg PetCO_2_ (post-hoc, t_(252)_=3.95, p=0.0006) while the initial decrease at +5mmHg PetCO_2_ did not reach significance (post-hoc, t_(252)_=0.6, p=0.93). No significant change of *PI*^*SE*^ with +5mmHg PetCO_2_ (post-hoc, t_(252)_=0.56, p=0.95) or +10mmHg PetCO_2_ could be observed (post-hoc, t_(252)_=1.70, p=0.32). PetO_2_ did not explain significant *PI*^*GE*/*SE*^ variance within-subject (Model 1: 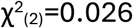, p=0.99) or between-subject (Model 1: 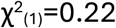, p=0.78). Changes in *PI*^*GE*/*SE*^ with hypercapnia were driven by *PA*^*GE*/*SE*^ (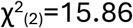, p=0.0007, Figure 4, second row), whereas the mean cardiac signal remained unchanged (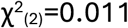, p=0.99, Figure 4, third row).

The cortical depth dependence of the pulsatility change with hypercapnia, 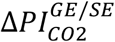, can be observed in Figure 5. The mean slope (A) and intercept (B) from the voxel-wise regression between *PI*^*GE*/*SE*^ and PetCO_2_ are displayed.

**Figure 5:**
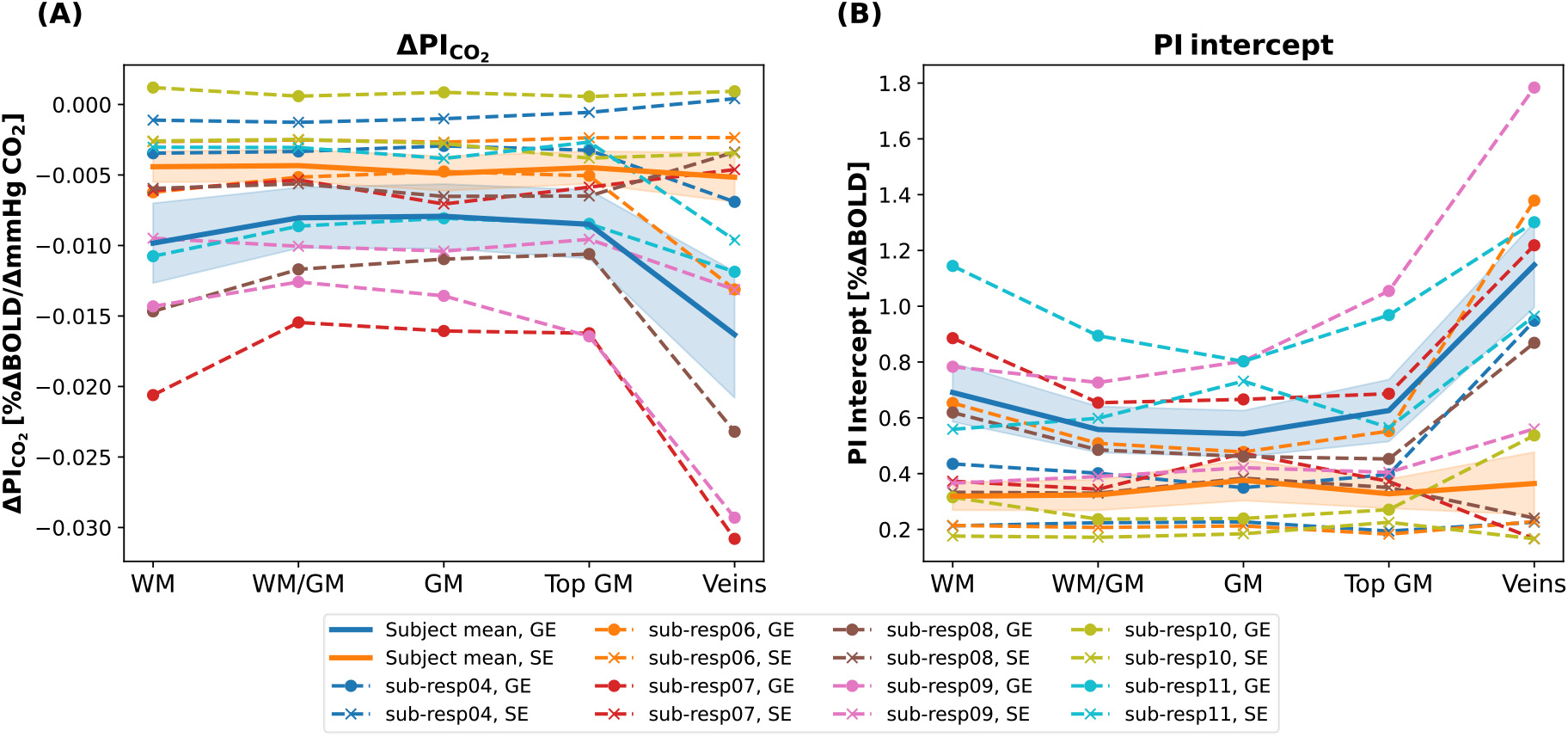
Change of Pulsatility Index (PI) with CO_2_ (ΔPI_CO2_) across cortical depth. ΔPI_CO2_ (A) and the intercept (B) are plotted for the individual subjects (dashed lines) and the subject average (thick lines, shaded are represents standard error of mean) for GE and SE.

The intercepts closely resemble baseline pulsatility curves (compare to Figure 4A), while negative values of the slope 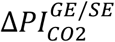 (the rate of change of *PI*^*GE*/*SE*^ with increasing PetCO_2_) indicate decreased *PI*^*GE*/*SE*^ with hypercapnia. The decrease differed by scan sequence (Model 2: 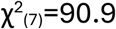, p=.0001) and laminae (Model 2: 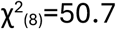, p<.001), with a significant interaction of the former indicating different spatial dependence of 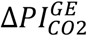 and 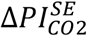 (Model 2: 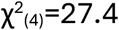, p=.0002). In GE, 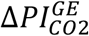 was significantly larger in veins than in low GM (post-hoc, t_(48)_=6.9, p<.000s1), middle GM (post-hoc, t_(48)_=7.0, p<.0001), top GM (post-hoc, t_(48)_=6.5, p<.0001), and WM (post-hoc, t_(48)_=5.4, p<.0001). No significant differences between other laminae were found. For SE, no significant differences between laminae were found for 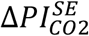. In summary, these results suggest that pulsatility decreases equally across cortical depth and only expresses a significantly larger decrease in large veins assessed by GE. The full linear mixed model coefficients and PBtest values for 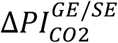 can be found in S9.

### 3.4 Pulsatility Index versus CVR and CBV

Figure 6 displays the linear relationship of baseline *PI*^*GE*/*SE*^ versus *CVR*^*GE*/*SE*^ (Figure 6A/C) and 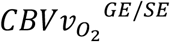 (Figure 6B/D), for all voxels responding significantly to hypercapnia and hyperoxia and excluding negative 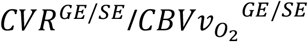-voxels.

**Figure 6:**
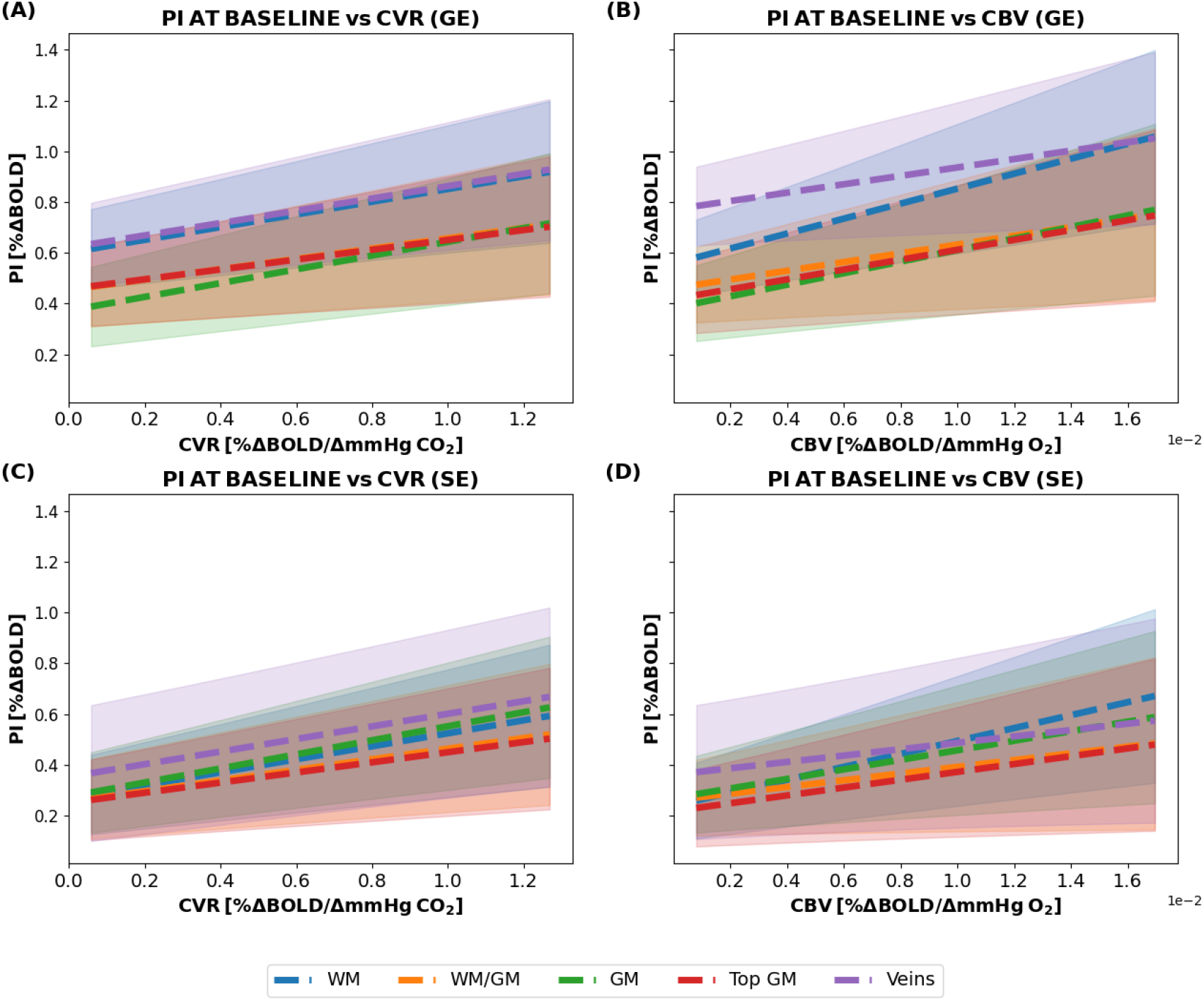
Pulsatility Index (PI) at Baseline versus CVR^GE^ (A), CVR^SE^ (C), 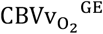 (B) and 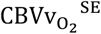 (D) as estimated by the Linear Mixed Model for the separate scan sequences and laminae. The shaded area represents the standard error of the mean for the population-level (fixed-effects) predictions computed using the variance–covariance matrix of the fixed effects.

To limit the abscissa range, plots were restricted to the 5–95th percentile *CVR*^*GE*/*SE*^ or 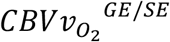 range, respectively. Mean *CVR*^*GE*/*SE*^ and 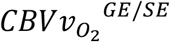 across cortical depth are reported in S1. *PI*^*GE*/*SE*^ correlated positively with *CVR*^*GE*/*SE*^ (Model 3: 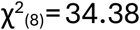, p=.0006) and *PI* changes with *CVR*^*GE*/*SE*^ were comparable for GE and SE (Model 3: 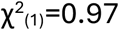, p=0.4). Similarly, a positive correlation between *PI*^*GE*/*SE*^ and 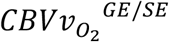 was observed (Model 4: 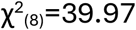, p=.0002), with interaction of scan sequence and 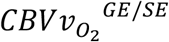 (Model 4: 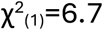, p=0.026) explaining further variance, indicating different *PI* changes with 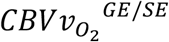 between the scan sequences.

Figure 7 shows a negative correlation of 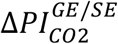 against 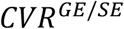 and 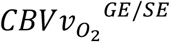, respectively.

**Figure 7:**
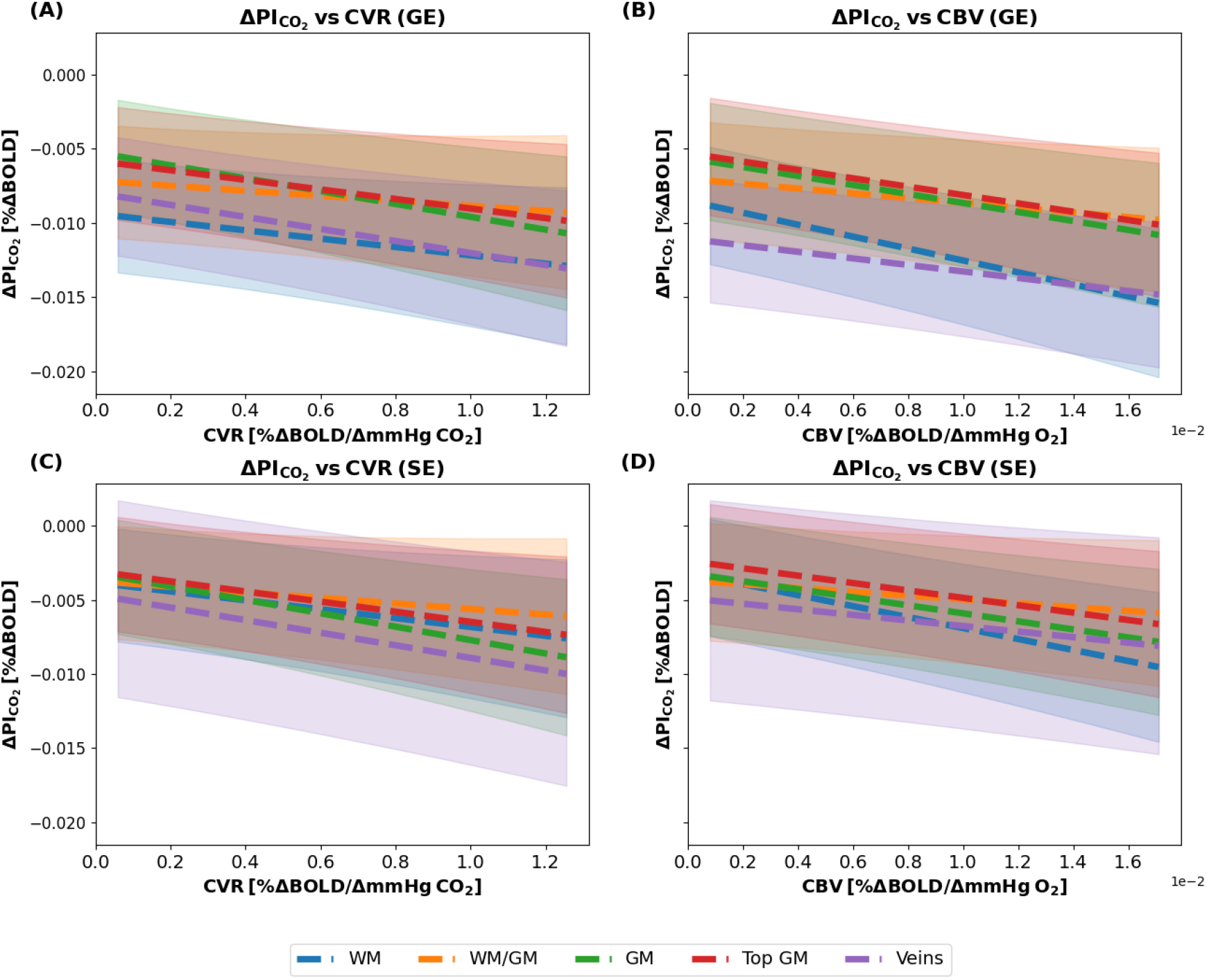
Change of Pulsatility Index with CO_2_ (ΔPI_CO2_) versus CVR^GE^ (A), CVR^SE^ (C), 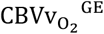 (B), and 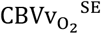 (D as estimated by the Linear Mixed Model for the separate scan sequences and laminae. The shaded area represents the standard error of the mean for the population-level (fixed-effects) predictions computed using the variance– covariance matrix of the fixed effects.

Significant variance of 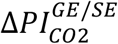 was explained by scan sequence (Model 5: 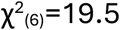, p=0.02) and *CVR*^*GE*/*SE*^ (Model 5: 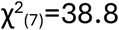, p=.0002), and scan sequence (Model 6: 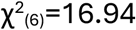, p=0.039) and 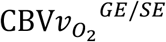 (Model 6: 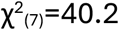, p<.0001).

All linear mixed model coefficients and PBtest tables for the above results are presented in the supplementary material (S10, S11, S12, S13).

## 4. Discussion

The aim of this study was to characterize the dependence of BOLD fMRI–based *PI* on vessel size, vascular density, and changes in vascular physiology. We derived pulsatility maps demonstrating physiologically plausible distributions of *PI*^*GE*/*SE*^, with increased values near pial veins. Pulsatility is larger in the macrovasculature as measured with *PI*^*GE*^ (sensitive to macro- and microvasculature) than in the microvasculature measured by *PI*^*SE*^ (selectively sensitive to the microvasculature) and the distinct laminar profiles of *PI*^*GE*/*SE*^ are consistent with the distribution of macro- and microvessels over cortical depth. The macrovascular dominated *PI*^*GE*^ decreased significantly during hypercapnia, demonstrating sensitivity to vascular tone, while the microvascular-dominated *PI*^*SE*^ did not change significantly.

### 4.1 Physiological Origin and Specificity of the BOLD-based Pulsatility Index

The BOLD fMRI-based *PI* is sensitive to blood volume and the deoxygenated hemoglobin concentration in blood, and therefore predominantly to capillaries and venous vessels (Ogawa et al., 1990). Additionally, BOLD fMRI is sensitive to flow effects (Bianciardi et al., 2016). Therefore, the *PI* may incorporate contributions of blood velocity and volume pulsations. In this analysis, low frequency oscillations were excluded and only changes around the cardiac frequencies are retained.

One possible source of capillary and venous volume and velocity pulsations are intravascular blood pressure pulsations. Pressure pulsations originate from the cardiac influx and propagate through the vascular system. Notably, pulsations of velocity and volume do not coincide. Rather, given that the flow through a vessel is computed as the product of lumen area and flow velocity, velocity and area pulsations are expected to be inversely proportional (van Tuijl et al., 2020). Against common perception, velocity and pressure pulsations persist in the capillaries (Bedggood & Metha, 2021; Rashid et al., 2012) and on the venous side (Bourquin et al., 2022; Driver et al., 2020). Therefore, it seems plausible that the pulsatile signal changes originate partially from capillary and venous volume and velocity pulsations.

Another mechanism that may induce venous cardiac pulsatility is venous compression. It occurs in large veins at the pial surface owing to intracranial pressure (ICP) oscillations. This mechanism hypothesizes that – following the Monro-Kellie Doctrine – brain-wide intracranial pressure pulsations cause the compression up to partial collapse of large veins as the intracranial pressure exceeds intravenous pressure. As a consequence, resistance to venous outflow increases. Intravenous pressure builds, and when it surpasses the intracranial pressure (ICP), flow is reestablished—resulting in a pulsatile character of venous outflow and intravenous pressure (De Simone et al., 2017). In addition to compression of superficial large veins, local venous compression inside brain tissue has been proposed as a driver of venous pulsatility. By inducing local tissue displacement, swelling arteries may compress adjacent venules, causing cardiac-frequent signal changes (Van Hulst et al., 2024).

### 4.2 Spatial Distribution and Vessel Size Dependence of the Pulsatility Index at Baseline

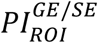 -estimates computed from ROI-average time-series yielded consistent laminae distributions and *PI*-values, highlighting robustness of the approach (compare Figure 4 and S2). The computed *PI*^*GE*/*SE*^ maps and cortical depth analysis show lower *PI*^*SE*^ than *PI*^*GE*^, consistent with lower microvascular pulsatility captured by SE due to its microvascular sensitivity. This likely reflects visco-elastic damping of arterial wall motion, dissipating pulsatile energy in large vessels and leading to reduced downstream microvascular pulsatility (Vikner et al., 2020; Zarrinkoob et al., 2016). In contrast, *PI*^*GE*^ are larger, reflecting macrovascular contributions where venous blood volume at lower pressure permits larger volume pulsations, while faster flow enhances flow effects. Our *PI*^*GE*^ matches the range observed by Chen et al. using retrospectively gated velocity selective ASL sensitive to all microvessels: in the GM they observe *PI* values of 0.38–0.68 (Chen et al., 2024).

The expected vascular specificity of our *PI*^*GE*/*SE*^ is supported by the spatial distribution of *PI*^*GE*/*SE*^ which aligns with vascular anatomy (Figure 3), with high *PI*^*GE*/*SE*^ near the pial surface and bordering the sagittal sinus, and proximal to large veins outlined by the vein mask. Venous compression provides a further explanation for larger *PI*^*GE*/*SE*^ proximal to veins: downstream veins carrying lower pressure, particularly those exposed to CSF at the cortical surface, are more easily compressed by tissue displacement and intracranial pressure fluctuations (De Simone et al., 2017). Unlike *PI*^*SE*^, *PI*^*GE*^ also show a trend for elevated WM pulsatility. This relates to a slight mean *PA*^*GE*^ increase toward WM, but not *PA*^*SE*^ (Figure 4), suggesting an underlying structure the GE BOLD-signal is sensitive to. Such may be veins in the WM with increased pulsatility or pulsating CSF-filled perivascular spaces (Lynch et al., 2023; Taoka et al., 2017). Notably, the low mean WM BOLD signal further amplifies *PI*^*GE*^ following the normalization, suggesting a methodological contribution to the observed increase. In contrast, *PI*^*SE*^ does not show this increase, possibly due to lower microvascular density in the WM. Further, capillaries may be subject to less compression, owing to higher intravascular pressure in capillaries, less tissue displacement due to smaller surrounding arterioles in the capillary bed, and greater wall stiffness from a higher thickness-to-diameter ratio (Müller et al., 2008).

A detailed comparison of *PI*^*GE*/*SE*^ was enabled by the cortical depth analysis. *PI*^*GE*^ decrease from WM into GM, then rise again toward the pial surface, and peak in larger pial veins. Contrary, for *PI*^*SE*^ no clear layer differences were identified post-hoc. Guo et al. reported decreasing CBV pulsatility with cortical depth using VASO (Guo et al., 2025). Compared to their *PI*, our *PI*^*GE*^ showed higher amplitudes (up to three times higher) and higher WM pulsatility, though both revealed increasing *PI* toward the cortical surface. Notably, VASO-MRI assesses predominantly arterial microvasculature, limiting comparability. Equally, Makedonov et al. and Theyers et al. found higher BOLD-based pulsatility in GM than WM (Makedonov et al., 2013; Theyers et al., 2019), while Adams and Sloots et al. reported larger volumetric strain in GM using DENSE-MRI, relating to increased volumetric pulsatility (Adams et al., 2020; Sloots et al., 2021). Contrary to *PI*^*GE*^, mean *PI*^*SE*^ peaks in mid-GM and declines toward both WM and the pial surface in our study. Notably, this effect did not reach significance. Nevertheless, Rashid et al. observed increased velocity *PI* in deeper GM capillaries in the rat brain, comparable to the peak of *PI*^*SE*^ observed here (Rashid et al., 2012). They attributed the increase to reduced compliance due to the absence of CSF and large draining veins in deeper tissue, leading to reduced vessel dilation and increased velocity *PI* ((van Tuijl et al., 2020)). However, while Rashid measured velocity *PI* from red blood cell tracking, BOLD *PI*^*SE*^ is also sensitive to volume changes, which should decline with increasing cortical depth if compliance and intravascular pulse pressure decreases. This discrepancy highlights the need to clarify relations between velocity- and volume-based *PI* and their respective roles in the BOLD signal.

### 4.3 Implications of PI correlating with CVR and CBV

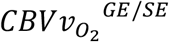 measures the voxel volume fraction occupied by capillary and venous vessels and varies significantly with cortical depth (compare S1: CVR, CBV across cortical depth, (Roefs et al., 2024)). Higher 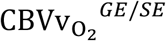 is associated with higher *PI* at baseline for GE and SE. 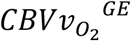 decreases significantly with depth, indicating that large veins dominate higher values (Schellekens et al., 2023). Accordingly, elevated *PI*^*GE*^ may arise from stronger flow effects, and/or more pulsatile venous volume. The linear correlation between *PI*^*SE*^ and 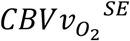 suggests higher *PI*^*SE*^ with greater microvascular density. However, the cortical depth profile of 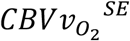 does not show the expected mid-GM peak found in prior studies (Guo et al., 2025; Zhao et al., 2006), likely due to confounding larger-vein signal or partial volume effects at the achieved resolution. This warrants further investigation of *PI*^*SE*^ - density relationships.

Additionally, *PI*^*GE*/*SE*^ correlates positively with *CVR*^*GE*/*SE*^, indicating that vessels with greater vasodilatory capacity also exhibit higher *PI*. Similarly, this correlation implies larger *PI* in large veins.

### 4.4 Response of Pulsatility Index to Vascular Changes

During hypercapnia, the *PI*^*GE*^ levels decrease. The visually apparent decrease of *PI*^*SE*^ on the other hand, is not significant (Figure 4). The *PI*^*GE*^ decrease is consistent across laminae and only significantly stronger in the large veins. As evident from the plots of *PA*^*GE*^ and mean cardiac signal, the decrease in *PI*^*GE*^ is driven by a decrease in the *PA*^*GE*^ (compare Figure 4, S7, S8). Previous studies’ findings raise the possibility of both reduced volume and velocity pulsations during hypercapnia (Casaccia et al., 2014; Guldbrandsen et al., 2024; Hachlouf et al., 2025; Hsu et al., 2004; Kucewicz et al., 2008; Naessens et al., 2023; Theyers et al., 2019; Van Bel et al., 1988). However, as it has been observed that velocity and volume pulsations are inversely related in larger vessels, a simultaneous lowering of both may not be feasible (van Tuijl et al., 2020). The following two paragraphs summarize evidence for decreased volume pulsations and velocity pulsations in response to hypercapnia, respectively. Overall, the studies concluding that hypercapnia lowers velocity *PI* outweigh arguments for lowered volume *PI*, suggesting that the hypercapnic decrease of *PI*^*GE*^ assessed by BOLD fMRI is dominated by changes in flow velocity. While our results do not provide a definite conclusion on the physiological underpinning of the observed *PI*^*GE*^ changes with hypercapnia, we can confirm that our methodology is able to detect subtle changes in vascular tone due to hypercapnia, proving the methods sensitivity to altered vascular physiology.

Vessel wall mechanical changes following hypercapnia-induced vasodilation imply reduced (relative) volume pulsatility during hypercapnia, as blood vessels exhibit a nonlinear stress– strain relationship, meaning their stiffness increases with dilation (Avolio et al., 2022). Accordingly, Casaccia et al. found that acute hypercapnia during rebreathing increased mean arterial pressure and carotid PWV (Casaccia et al., 2014), a measure of vascular stiffness (Townsend, 2022). Further, following vascular dilation during hypercapnia, the cerebral compliance reduces due to a preceding compression of compliant structures, and ICP would increase acting against vascular volume pulsations (Piper Ian et al., 1992; Wagshul et al., 2011b). The two described mechanisms reduce vascular distensibility in response to pressure pulsations. Further, these findings are in line with observations made by Naessens et al, who observed a reduced amplitude of ICA diameter pulsations in systemic hypertensive rats, where hypertension is assumed to induce macrovascular vasodilation as well (Naessens et al., 2023). Notably, however, the longer time-scale of chronic hypertension may induce structural changes beyond the scope of hypercapnia-induced changes. We cannot conclude whether the stiffening of vessel walls or reduced cerebral compliance dominates *PI*^*GE*^ changes during hypercapnia based on our observations. Nevertheless, reduced vessel wall distensibility could result in a lower volume-based *PI*^*GE*^.

On the other hand, several studies examined pulsatility during hypercapnia and found consistently reduced velocity *PI* in large arteries (MCA, ACA), observed across healthy adults, cardiac arrest patients, and infants (Guldbrandsen et al., 2024; Hachlouf et al., 2025; Hsu et al., 2004; Robertson et al., 2008; Van Bel et al., 1988). Acetazolamide-induced vasodilation produced similar reductions alongside increased intracranial pressure (Hachlouf et al., 2025), and animal models confirmed decreased hypercapnic velocity *PI* and vascular resistance (Czosnyka et al., 1996). Plethysmography-based tissue pulsatility imaging found decreasing pulse amplitudes with decreasing end-tidal CO_2_ (Kucewicz et al., 2008). These results capture tissue deformation which more likely coincides with blood volume pulsations, and therefore align with decreased velocity *PI* upon hypercapnia. The comparability to our *PI*^*GE*/*SE*^ is however limited as no signal normalization was conducted. At the microvascular level, diffuse correlation spectroscopy (DCS) revealed decreased *PI* during 5% CO_2_ inhalation, despite elevated blood pressure, with sex differences at baseline (Urner et al., 2023; Zhou et al., 2021). DCS predominantly measures flow velocity, therefore reflecting a velocity *PI* (Wang et al., 2024). The decrease of velocity *PI* during hypercapnia is supported by our observation that *PI*^*GE*^, sensitive to larger vessels with higher flow velocities, decreases significantly during hypercapnia, while the decreasing trend does not reach significance for *PI*^*SE*^, reflecting predominantly capillary vessels with low flow velocities. Similarly, the negative correlation of 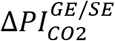 to *CVR*^*GE*/*SE*^ and 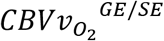 implies that vessels with a stronger response to hypercapnia and/or larger baseline blood volume, commonly larger vessels located at the pial surface (compare S1), experience a stronger decrease in *PI* in response to hypercapnia.

Finally, a previous study indicates that the heart rate affects observed volume pulsatility (Naessens et al., 2023). We did not see significantly elevated heart rates in six out of seven subjects during hypercapnia (compare S4: Cardiac Beats Per Minute), suggesting that the observed *PI*^*GE*^ reduction is more likely attributable to mechanical changes in the vasculature.

### 4.5 Methodological Contributions and Clinical Relevance

Compared to specialized MRI sequences such as DENSE, VASO, or VS-ASL, our method relies on standard fMRI sequences, provides high spatial resolution, and enables retrospective analysis of existing datasets from clinical populations. Cardiac traces can be extracted from fMRI data with existing software packages, eliminating the need for a measured cardiac trace (Aslan et al., 2019). Given that *PI* alterations have been associated with various cerebrovascular and neurodegenerative diseases (Delli Pizzi et al., 2023; Roher et al., 2011; Shi et al., 2018, 2020), and that early pathological changes are expected to originate in the microvasculature (Van De Haar et al., 2016), there is a need for non-invasive tools capable of resolving pulsatility at this scale. Our findings contribute to a better understanding of the specificity of BOLD-based *PI* to different vascular compartments, an essential step toward interpreting pulsatility alterations in vascular pathologies.

Several MRI approaches have been used to probe pulsatility in the brain. DENSE-MRI can quantify cardiac-induced volumetric strain from captured tissue motion (Sloots et al., 2021). While DENSE-derived volumetric strain is predominantly selectively specific to vascular volume pulsations, it has limited resolution which hampers analysis across cortical depth. It also lacks vessel-size specificity, similar to standard GE-based BOLD. Guo et al. used retrospectively gated VASO-MRI to map CBV pulsations across cortical depth (Guo et al., 2025). Their *PI*, derived as pulse amplitude normalized by mean CBV, presumably reflects volume pulsations from arteries to venules, though inflow effects may contribute. They demonstrated depth-dependent changes but could not separate macro- from microvascular signals. Chen et al. applied retrospectively gated velocity-selective arterial spin labeling (VS-ASL) to develop an arterial flow-weighted *PI* (Chen et al., 2024). Like Guo, they used normalized signal extrema over the cardiac cycle, but did not assess depth-dependent changes. Similar to our study, they did not achieve a clear distinction between volume and velocity pulsatility. Other studies quantified normalized variance of rs-fMRI time series as proxy for pulsatility, but without vascular or depth specificity (Makedonov et al., 2013; Shirzadi et al., 2018; Tuovinen et al., 2020). Some assessed cardiac spectral power, requiring high temporal sampling to avoid aliasing, thereby reducing field of view or spatial resolution (Makedonov et al., 2013; Viessmann et al., 2019). The required temporal resolution was often addressed by retrospective gating or deep learning to reconstruct cardiac signals (Atwi et al., 2020; Kim et al., 2021; Potvin-Jutras et al., 2025; Theyers et al., 2019; Valsamis et al., 2023; Viessmann et al., 2017). Overall, no method has fully linked pulsatility to the underlying vascular anatomy and physiology. Our method similarly used a small field of view and retrospective gating to increase temporal resolution. Combining SE and GE readouts at 7T enabled separation of macro- and microvascular contributions and cortical-depth mapping. By relating pulsatility to *CVR*^*GE*/*SE*^ and venous 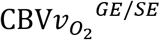 measures, we further linked observed pulsatility to microvascular physiology.

### 4.6 Limitations and Future Work

Our study has several limitations that should be acknowledged. The requirement of a PPU signal adds complication to the data acquisition and limits retrospective applicability. There were visual stimuli present in our data which were excluded by treating the corresponding hemodynamics response functions as nuisance regressors. Further, we only used data acquired at a single TR with a FOV covering the visual cortex. This precludes an understanding on the TR dependence of our methodology and limits the generalizability of our findings to all brain areas. Together with the small sample size these limitations warrant replication in larger cohorts at varying TRs and with larger FOVs.

Methodologically, our approach does not allow us to distinguish fully between pulsatility arising from changes in vascular volume versus flow velocity, both of which can contribute to BOLD signal fluctuations. Finally, like most studies of cerebral pulsatility, we are unable to disentangle contributions from various drivers, such as intravascular pressure, extravascular factors such as ICP fluctuations or tissue displacement, and vessel wall properties (compliance).

## 5. Conclusion

This study advances the interpretability of BOLD-based pulsatility indices (*PI*) by systematically characterizing their dependence on vessel size, compartment, and physiological state. Using both GE and SE BOLD acquisitions at 7T, we disentangled macro- and microvascular contributions and revealed distinct *PI*^*GE*/*SE*^ profiles across cortical depth. *PI*^*GE*^ were overall larger than *PI*^*SE*^ and higher *PI*^*GE*^ were observed near large pial veins and in white matter, reflecting macrovascular sensitivity and venous pulsations. Positive correlations with venous blood volume and cerebrovascular reactivity suggest that both vascular density and vessel wall mechanics influence *PI*^*GE*/*SE*^, highlighting that *PI*^*GE*/*SE*^ is not a pure marker of vascular resistance. The sensitivity of *PI*^*GE*^ to physiological modulation as a proxy for pathological changes was demonstrated by altered *PI*^*GE*^ under hypercapnia. *PI*^*SE*^, on the other hand, are less affected by changes in vascular tone. As the method relies only on standard fMRI sequences, it is well-suited for retrospective application and potentially translatable to lower clinical field strengths. Together, these findings support the utility of BOLD-based *PI*^*GE*/*SE*^ as a spatially and physiologically informative measure of cerebrovascular health.

## 6. Data and Code availability

All data and code will be shared upon reasonable request.

## 7. Author Contributions

Hans Christian Rundfeldt conceived and designed the study, performed all analyses, and wrote the manuscript. Wouter Schellekens contributed previously acquired data. Emiel C.A. Roefs assisted with data acquisition and provided code used in the study. Alex A. Bhogal supported data acquisition. Mario Gilberto Báez-Yáñez, Jaco J.M. Zwanenburg, and Natalia Petridou contributed to project conceptualization and discussions and provided institutional support. All authors reviewed and approved the final manuscript.

## 8. Funding

The study was supported by the Netherlands organization for health research and development (ZonMw) under award number 10510032120006 (MODEM program), Alzheimer Netherlands under award number WE.30-2022-04, and the National Institute of Mental Health of the National Institutes of Health under the Award Number R01MH111417. The content is solely the responsibility of the authors and does not necessarily represent the official views of the National Institutes of Health.

## 9. Declaration of Competing Interests

The author(s) declared no potential conflicts of interest with respect to the research, authorship, and/or publication of this article.

## 10. Acknowledgments

We gratefully acknowledge the support and discussions with the *Mechanisms Of DEMentia* (MODEM) and *Cerebral HemodynamIcs, Metabolism and clearancE* (CHIME) PhD consortiums. This publication is part of the MODEM consortium and project CHIME (file number P22.012) of the research programme Perspectief, which is (partly) financed by the Dutch Research Council (NWO).

## Notes

### Competing Interest Statement

The authors have declared no competing interest.

## References

Aaslid, R., Markwalder, T. M., & Nornes, H. (1982). Noninvasive transcranial Doppler ultrasound recording of flow velocity in basal cerebral arteries. Journal of Neurosurgery, 57(6), 769–774. 10.3171/JNS.1982.57.6.0769

Adams, A. L., Viergever, M. A., Luijten, P. R., & Zwanenburg, J. J. M. (2020). Validating faster DENSE measurements of cardiac-induced brain tissue expansion as a potential tool for investigating cerebral microvascular pulsations. NeuroImage, 208. 10.1016/j.neuroimage.2019.116466

Arts, T., Onkenhout, L. P., Amier, R. P., van der Geest, R., van Harten, T., Kappelle, J., Kuipers, S., van Osch, M. J. P., van Bavel, E. T., Biessels, G. J., & Zwanenburg, J. J. M. (2022). Non-Invasive Assessment of Damping of Blood Flow Velocity Pulsatility in Cerebral Arteries With MRI. Journal of Magnetic Resonance Imaging, 55(6), 1785–1794. 10.1002/JMRI.27989

Aslan, S., Hocke, L., Schwarz, N., & Frederick, B. (2019). Extraction of the cardiac waveform from simultaneous multislice fMRI data using slice sorted averaging and a deep learning reconstruction filter. NeuroImage, 198, 303–316. 10.1016/J.NEUROIMAGE.2019.05.049

Atwi, S., Robertson, A. D., Theyers, A. E., Ramirez, J., Swartz, R. H., Marzolini, S., & MacIntosh, B. J. (2020). Cardiac-Related Pulsatility in the Insula Is Directly Associated With Middle Cerebral Artery Pulsatility Index. Journal of Magnetic Resonance Imaging, 51(5), 1454–1462. 10.1002/JMRI.26950

Avolio, A., Spronck, B., Tan, I., Cox, J., & Butlin, M. (2022). Basic principles that determine relationships between pulsatile hemodynamic phenomena and function of elastic vessels. In J. A. Chirinos (Ed.), Textbook of arterial stiffness and pulsatile hemodynamics in health and disease (1st ed., pp. 3–26). Academic Press.

Balduzzi, D., Riedner, B. A., & Tononi, G. (2008). A BOLD window into brain waves. Proceedings of the National Academy of Sciences, 105(41), 15641–15642. 10.1073/PNAS.0808310105

Bedggood, P., & Metha, A. (2021). Direct measurement of pulse wave propagation in capillaries of the human retina. Optics Letters, Vol. 46, Issue 18, Pp. 4450-4453, 46(18), 4450–4453. 10.1364/OL.434454

Bernier, M., Cunnane, S. C., & Whittingstall, K. (2018). The morphology of the human cerebrovascular system. Human Brain Mapping, 39(12), 4962–4975. 10.1002/HBM.24337

Bhogal, A. A., De Vis, J. B., Siero, J. C. W., Petersen, E. T., Luijten, P. R., Hendrikse, J., Philippens, M. E. P., & Hoogduin, H. (2016). The BOLD cerebrovascular reactivity response to progressive hypercapnia in young and elderly. NeuroImage, 139, 94–102. 10.1016/j.neuroimage.2016.06.010

Bianciardi, M., Toschi, N., Polimeni, J. R., Evans, K. C., Bhat, H., Keil, B., Rosen, B. R., Boas, D. A., & Wald, L. L. (2016). The pulsatility volume index: an indicator of cerebrovascular compliance based on fast magnetic resonance imaging of cardiac and respiratory pulsatility. Philosophical Transactions of the Royal Society A: Mathematical, Physical and Engineering Sciences, 374(2067). 10.1098/RSTA.2015.0184

Birn, R. M., Smith, M. A., Jones, T. B., & Bandettini, P. A. (2008). The respiration response function: the temporal dynamics of fMRI signal fluctuations related to changes in respiration. NeuroImage, 40(2), 644–654. 10.1016/J.NEUROIMAGE.2007.11.059

Bourquin, C., Poree, J., Lesage, F., & Provost, J. (2022). In Vivo Pulsatility Measurement of Cerebral Microcirculation in Rodents Using Dynamic Ultrasound Localization Microscopy. IEEE Transactions on Medical Imaging, 41(4), 782–792. 10.1109/TMI.2021.3123912

Budde, J., Shajan, G., Zaitsev, M., Scheffler, K., & Pohmann, R. (2014). Functional MRI in human subjects with gradient-echo and spin-echo EPI at 9.4 T. Magnetic Resonance in Medicine, 71(1), 209–218. 10.1002/MRM.24656

Casaccia, S., Sirevaag, E. J., Richter, E., O’Sullivan, J. A., Scalise, L., & Rohrbaugh, J. W. (2014). Decoding carotid pressure waveforms recorded by laser Doppler vibrometry: Effects of rebreathing. AIP Conference Proceedings, 1600, 298–312. 10.1063/1.4879596

Chen, C., Barnes, R. A., Bangen, K. J., Han, F., Pfeuffer, J., Wong, E. C., Liu, T. T., & Bolar, D. S. (2024). MVP-VSASL: measuring MicroVascular Pulsatility using velocity-selective arterial spin labeling. Magnetic Resonance in Medicine. 10.1002/MRM.30370

Chiarelli, P. A., Bulte, D. P., Wise, R., Gallichan, D., & Jezzard, P. (2007). A calibration method for quantitative BOLD fMRI based on hyperoxia. NeuroImage, 37(3), 808–820. 10.1016/j.neuroimage.2007.05.033

Czosnyka, M., Richards, H. K., Whitehouse, H. E., & Pickard, J. D. (1996). Relationship between transcranial Doppler-determined pulsatility index and cerebrovascular resistance: an experimental study. Journal of Neurosurgery, 84(1), 79–84. 10.3171/JNS.1996.84.1.0079

De Simone, R., Ranieri, A., & Bonavita, V. (2017). Starling resistors, autoregulation of cerebral perfusion and the pathogenesis of idiopathic intracranial hypertension. In Panminerva Medica (Vol. 59, Number 1, pp. 76–89). Edizioni Minerva Medica. 10.23736/S0031-0808.16.03248-1

Delli Pizzi, S., Gambi, F., Di Pietro, M., Caulo, M., Sensi, S. L., & Ferretti, A. (2023). BOLD cardiorespiratory pulsatility in the brain: from noise to signal of interest. Frontiers in Human Neuroscience, 17, 1327276. 10.3389/FNHUM.2023.1327276/BIBTEX

Driver, I. D., Traat, M., Fasano, F., & Wise, R. G. (2020). Most Small Cerebral Cortical Veins Demonstrate Significant Flow Pulsatility: A Human Phase Contrast MRI Study at 7T. Frontiers in Neuroscience, 14. 10.3389/fnins.2020.00415

Duong, T. Q., Yacoub, E., Adriany, G., Hu, X., Uğurbil, K., & Kim, S. G. (2003). Microvascular BOLD contribution at 4 and 7 T in the human brain: Gradient-echo and spin-echo fMRI with suppression of blood effects. Magnetic Resonance in Medicine, 49(6), 1019–1027. 10.1002/MRM.10472

Glover, G. H., Li, T.-Q., & Ress, D. (2000). Image-Based Method for Retrospective Correction of Physiological Motion Effects in fMRI: RETROICOR. 10.1002/1522-2594(200007)44:1

Guldbrandsen, H., Juhl-Olsen, P., Eastwood, G. M., Wethelund, K. L., & Grejs, A. M. (2024). Sonographic evaluation of intracranial hemodynamics and pressure after out-of-hospital cardiac arrest: An exploratory sub-study of the TAME trial. Critical Care and Resuscitation, 26(3), 176–184. 10.1016/J.CCRJ.2024.06.001

Guo, F., Zhao, C., Shou, Q., Jin, N., Jann, K., Shao, X., & Wang, D. J. (2025). Assessing Cerebral Microvascular Volumetric Pulsatility with High-Resolution 4D CBV MRI at 7T. Nat Cardiovasc Res, 1424–1438. 10.1038/s44161-025-00722-1

Hachlouf, A., Stella, C., Cavalli, I., Gouvêa Bogossian, E., Schuind, S., Anderloni, M., & Taccone, F. S. (2025). Effects of acetazolamide on intracranial pressure and brain tissue oxygenation on patients with acute brain injury: A pilot physiological study. Physiological Reports, 13(1), e70159. 10.14814/PHY2.70159

Halekoh, U., & Højsgaard, S. (2014). A Kenward-Roger Approximation and Parametric Bootstrap Methods for Tests in Linear Mixed Models – The R Package pbkrtest. Journal of Statistical Software, 59(9), 1–32. 10.18637/JSS.V059.I09

Hsu, H. Y., Chern, C. M., Kuo, J. S., Kuo, T. B. J., Chen, Y. T., & Hu, H. H. (2004). Correlations among critical closing pressure, pulsatility index and cerebrovascular resistance. Ultrasound in Medicine & Biology, 30(10), 1329–1335. 10.1016/J.ULTRASMEDBIO.2004.08.006

Huber, L. (Renzo) R., Poser, B. A., Bandettini, P. A., Arora, K., Wagstyl, K., Cho, S., Goense, J., Nothnagel, N., Morgan, A. T., van den Hurk, J., Müller, A. K., Reynolds, R. C., Glen, D. R., Goebel, R., & Gulban, O. F. (2021). LayNii: A software suite for layer-fMRI. NeuroImage, 237, 118091. 10.1016/J.NEUROIMAGE.2021.118091

Huntenburg, J. M., Steele, C. J., & Bazin, P. L. (2018). Nighres: processing tools for high-resolution neuroimaging. GigaScience, 7(7), 1–9. 10.1093/GIGASCIENCE/GIY082

Iliff, J. J., Wang, M., Zeppenfeld, D. M., Venkataraman, A., Plog, B. A., Liao, Y., Deane, R., & Nedergaard, M. (2013). Cerebral Arterial Pulsation Drives Paravascular CSF–Interstitial Fluid Exchange in the Murine Brain. Journal of Neuroscience, 33(46), 18190–18199. 10.1523/JNEUROSCI.1592-13.2013

Kim, T., Kim, S. Y., Agarwal, V., Cohen, A., Roush, R., Chang, Y. F., Cheng, Y., Snitz, B., Huppert, T. J., Bagic, A., Kamboh, M. I., Doman, J., & Becker, J. T. (2021). Cardiac-induced cerebral pulsatility, brain structure, and cognition in middle and older-aged adults. NeuroImage, 233, 117956. 10.1016/J.NEUROIMAGE.2021.117956

Kiviniemi, V., Wang, X., Korhonen, V., Keinänen, T., Tuovinen, T., Autio, J., Levan, P., Keilholz, S., Zang, Y. F., Hennig, J., & Nedergaard, M. (2016). Ultra-fast magnetic resonance encephalography of physiological brain activity-Glymphatic pulsation mechanisms? Journal of Cerebral Blood Flow and Metabolism, 36(6), 1033–1045. 10.1177/0271678X15622047/ASSET/IMAGES/LARGE/10.1177_0271678X15622047-FIG5.JPEG

Kornemann, N., Klimeš, F., Kern, A. L., Behrendt, L., Voskrebenzev, A., Gutberlet, M., Wattjes, M. P., Wacker, F., Vogel-Claussen, J., & Glandorf, J. (2023). Cerebral microcirculatory pulse wave propagation and pulse wave amplitude mapping in retrospectively gated MRI. Scientific Reports 2023 13:1, 13(1), 1–10. 10.1038/s41598-023-48439-0

Kucewicz, J. C., Dunmire, B., Giardino, N. D., Leotta, D. F., Paun, M., Dager, S. R., & Beach, K. W. (2008). Tissue Pulsatility Imaging of Cerebral Vasoreactivity During Hyperventilation. Ultrasound in Medicine & Biology, 34(8), 1200–1208. 10.1016/J.ULTRASMEDBIO.2008.01.001

Kuznetsova, A., Brockhoff, P. B., & Christensen, R. H. B. (2017). lmerTest Package: Tests in Linear Mixed Effects Models. Journal of Statistical Software, 82(13), 1–26. 10.18637/JSS.V082.I13

Lynch, K. M., Sepehrband, F., Toga, A. W., & Choupan, J. (2023). Brain perivascular space imaging across the human lifespan. NeuroImage, 271, 120009. 10.1016/J.NEUROIMAGE.2023.120009

Makedonov, I., Black, S. E., & MacIntosh, B. J. (2013). BOLD fMRI in the White Matter as a Marker of Aging and Small Vessel Disease. PLOS ONE, 8(7), e67652. 10.1371/JOURNAL.PONE.0067652

Mestre, H., Tithof, J., Du, T., Song, W., Peng, W., Sweeney, A. M., Olveda, G., Thomas, J. H., Nedergaard, M., & Kelley, D. H. (2018). Flow of cerebrospinal fluid is driven by arterial pulsations and is reduced in hypertension. Nature Communications 2018 9:1, 9(1), 1–9. 10.1038/s41467-018-07318-3

Mitchell, G. F. (2008). Effects of central arterial aging on the structure and function of the peripheral vasculature: implications for end-organ damage. Journal of Applied Physiology (Bethesda, Md. : 1985), 105(5), 1652–1660. 10.1152/JAPPLPHYSIOL.90549.2008

Müller, B., Lang, S., Dominietto, M., Rudin, M., Schulz, G., Deyhle, H., Germann, M., Pfeiffer, F., David, C., & Weitkamp, T. (2008). High-resolution tomographic imaging of microvessels. Developments in X-Ray Tomography VI, 7078, 70780B. 10.1117/12.794157

Naessens, D. M. P., de Vos, J., Richard, E., Wilhelmus, M. M. M., Jongenelen, C. A. M., Scholl, E. R., van der Wel, N. N., Heijst, J. A., Teunissen, C. E., Strijkers, G. J., Coolen, B. F., VanBavel, E., & Bakker, E. N. T. P. (2023). Effect of long-term antihypertensive treatment on cerebrovascular structure and function in hypertensive rats. Scientific Reports, 13(1), 3481. 10.1038/S41598-023-30515-0

Ogawa, S., Lee, T. M., Kay, A. R., & Tank, D. W. (1990). Brain magnetic resonance imaging with contrast dependent on blood oxygenation. Proceedings of the National Academy of Sciences, 87(24), 9868–9872. 10.1073/PNAS.87.24.9868

Panchuelo, R. M. S., Schluppeck, D., Harmer, J., Bowtell, R., & Francis, S. (2015). Assessing the Spatial Precision of SE and GE-BOLD Contrast at 7 Tesla. Brain Topography, 28(1), 62–65. 10.1007/S10548-014-0420-4/FIGURES/2

Pantoni, L. (2010). Cerebral small vessel disease: from pathogenesis and clinical characteristics to therapeutic challenges. The Lancet Neurology, 9(7), 689–701. 10.1016/S1474-4422(10)70104-6/ASSET/9BBFBA4D-3FBE-407B-BC7F-BEC6D85DAF29/MAIN.ASSETS/GR3.JPG

Penny, W., Friston, K., Ashburner, J., Kiebel, S., & Nichols, T. (2007). Statistical Parametric Mapping: The Analysis of Functional Brain Images. In Statistical Parametric Mapping: The Analysis of Functional Brain Images. Elsevier Ltd. 10.1016/B978-0-12-372560-8.X5000-1

Pham, S. D. T., Chatziantoniou, · C, Van Vliet, · J T, Van Tuijl, · R J, Bulk, · M, Costagli, · M, De Rochefort, · L, Kraff, · O, Ladd, · M E, Pine, · K, Ronen, · I, Siero, · J C W, Tosetti, · M, Villringer, · A, Biessels, · G J, & Zwanenburg, · J J M. (2025). Blood Flow Velocity Analysis in Cerebral Perforating Arteries on 7T 2D Phase Contrast MRI with an Open-Source Software Tool (SELMA). Neuroinformatics 2025 23:2, 23(2), 1–13. 10.1007/S12021-024-09703-4

Piper Ian, R., Chan, K. H., Whittle, I. R., & Miller, J. D. (1992). An Experimental Study of Cerebrovascular Resistance, Pressure Transmission, and Craniospinal Compliance. Neurosurgery, 32(5), 805–816. 10.1227/00006123-199305000-00014

Potvin-Jutras, Z., Intzandt, B., Mohammadi, H., Liu, P., Chen, J. J., & Gauthier, C. J. (2025). Sex-specific effects of intensity and dose of physical activity on BOLD-fMRI cerebrovascular reactivity and cerebral pulsatility. Journal of Cerebral Blood Flow and Metabolism. 10.1177/0271678X251325399

Pruim, R. H. R., Mennes, M., van Rooij, D., Llera, A., Buitelaar, J. K., & Beckmann, C. F. (2015). ICA-AROMA: A robust ICA-based strategy for removing motion artifacts from fMRI data. NeuroImage, 112, 267–277. 10.1016/J.NEUROIMAGE.2015.02.064

Rajna, Z., Mattila, H., Huotari, N., Tuovinen, T., Krüger, J., Holst, S. C., Korhonen, V., Remes, A. M., Seppänen, T., Hennig, J., Nedergaard, M., & Kiviniemi, V. (2021). Cardiovascular brain impulses in Alzheimer’s disease. Brain, 144(7), 2214–2226. 10.1093/BRAIN/AWAB144

Rashid, S., McAllister, J. P., Yu, Y., & Wagshul, M. E. (2012). Neocortical capillary flow pulsatility is not elevated in experimental communicating hydrocephalus. Journal of Cerebral Blood Flow and Metabolism, 32(2), 318–329. 10.1038/JCBFM.2011.130/ASSET/71F69AB5-DA03-4CCF-91C3-EDFCCE1959F2/ASSETS/IMAGES/LARGE/10.1038_JCBFM.2011.130-FIG7.JPG

research, R. C.-C. and B., & 1996, undefined. (1996). AFNI: software for analysis and visualization of functional magnetic resonance neuroimages. Elsevier, 29, 162–173. https://www.sciencedirect.com/science/article/pii/S0010480996900142

Robertson, J. W., Debert, C. T., Frayne, R., & Poulin, M. J. (2008). Variability of Middle Cerebral Artery Blood Flow with Hypercapnia in Women. Ultrasound in Medicine and Biology, 34(5), 730–740. 10.1016/j.ultrasmedbio.2007.07.024

Roefs, E. C. A., Schellekens, W., Báez-Yáñez, M. G., Bhogal, A. A., Groen, I. I. A., van Osch, M. J. P., Siero, J. C. W., & Petridou, N. (2024). The contribution of the vascular architecture and cerebrovascular reactivity to the BOLD signal formation across cortical depth. Imaging Neuroscience, 2, 1–19. 10.1162/IMAG_A_00203

Roher, A. E., Garami, Z., Tyas, S. L., Maarouf, C. L., Kokjohn, T. A., Belohlavek, M., Vedders, L. J., Connor, D., Sabbagh, M. N., Beach, T. G., & Emmerling, M. R. (2011). Transcranial Doppler ultrasound blood flow velocity and pulsatility index as systemic indicators for Alzheimer’s disease. Alzheimer’s & Dementia, 7(4), 445–455. 10.1016/J.JALZ.2010.09.002

Saad, Z. S., Glen, D. R., Chen, G., Beauchamp, M. S., Desai, R., & Cox, R. W. (2008). A New Method for Improving Functional-to-Structural MRI Alignment using Local Pearson Correlation. NeuroImage, 44(3), 839. 10.1016/J.NEUROIMAGE.2008.09.037

Schellekens, W., Bhogal, A. A., Roefs, E. C. A., Báez-Yáñez, M. G., Siero, J. C. W., & Petridou, N. (2023). The many layers of BOLD. The effect of hypercapnic and hyperoxic stimuli on macro- and micro-vascular compartments quantified by CVR, M, and CBV across cortical depth. Journal of Cerebral Blood Flow and Metabolism, 43(3), 419–432. 10.1177/0271678X221133972/ASSET/IMAGES/LARGE/10.1177_0271678X221133972-FIG7.JPEG

Shi, Y., Thrippleton, M. J., Blair, G. W., Dickie, D. A., Marshall, I., Hamilton, I., Doubal, F. N., Chappell, F., & Wardlaw, J. M. (2020). Small vessel disease is associated with altered cerebrovascular pulsatility but not resting cerebral blood flow. Journal of Cerebral Blood Flow and Metabolism, 40(1), 85–99. 10.1177/0271678X18803956/ASSET/IMAGES/LARGE/10.1177_0271678X18803956-FIG3.JPEG

Shi, Y., Thrippleton, M. J., Marshall, I., & Wardlaw, J. M. (2018). Intracranial pulsatility in patients with cerebral small vessel disease: a systematic review. Clinical Science, 132(1), 157–171. 10.1042/CS20171280

Shirzadi, Z., Robertson, A. D., Metcalfe, A. W., Duff-Canning, S., Marras, C., Lang, A. E., Masellis, M., & MacIntosh, B. J. (2018). Brain tissue pulsatility is related to clinical features of Parkinson’s disease. NeuroImage: Clinical, 20, 222–227. 10.1016/J.NICL.2018.07.017

Sleight, E. (2023). Optimisation, evaluation and application of cerebrovascular reactivity measurement using magnetic resonance imaging in patients with cerebral small vessel disease [PhD Thesis]. University of Edinburgh.

Slessarev, M., Han, J., Mardimae, A., Prisman, E., Preiss, D., Volgyesi, G., Ansel, C., Duffin, J., & Fisher, J. A. (2007). Prospective targeting and control of end-tidal CO2 and O2 concentrations. The Journal of Physiology, 581(3), 1207–1219. 10.1113/JPHYSIOL.2007.129395

Sloots, J. J., Biessels, G. J., de Luca, A., & Zwanenburg, J. J. M. (2021). Strain Tensor Imaging: Cardiac-induced brain tissue deformation in humans quantified with high-field MRI. NeuroImage, 236, 118078. 10.1016/J.NEUROIMAGE.2021.118078

Taoka, T., Fukusumi, A., Miyasaka, T., Kawai, H., Nakane, T., Kichikawa, K., & Naganawa, S. (2017). Structure of the medullary veins of the cerebral hemisphere and related disorders. Radiographics, 37(1), 281–297. 10.1148/RG.2017160061/ASSET/IMAGES/LARGE/RG.2017160061.FIG10B.JPEG

Theyers, A. E., Goldstein, B. I., Metcalfe, A. W. S., Robertson, A. D., & MacIntosh, B. J. (2019). Cerebrovascular blood oxygenation level dependent pulsatility at baseline and following acute exercise among healthy adolescents. Journal of Cerebral Blood Flow and Metabolism, 39(9), 1737–1749. 10.1177/0271678X18766771

Townsend, R. R. (2022). Arterial stiffness and pulsatile hemodynamics in renal disease. In J. Chirinos (Ed.), Textbook of arterial stiffness and pulsatile hemodynamics in health and disease (1st ed., pp. 637–647). Academic Press.

Tuovinen, T., Häkli, J., Rytty, R., Krüger, J., Korhonen, V., Järvelä, M., Helakari, H., Kananen, J., Nikkinen, J., Veijola, J., Remes, A. M., Kiviniemi, V., Rosen, H., Dickerson, B. C., Domoto-Reilly, K., Knopman, D., Boeve, B. F., Boxer, A. L., Kornak, J., … Mandelli, M. L. (2024). The relative brain signal variability increases in the behavioral variant of frontotemporal dementia and Alzheimer’s disease but not in schizophrenia. Journal of Cerebral Blood Flow and Metabolism. 10.1177/0271678X241262583

Tuovinen, T., Kananen, J., Rajna, Z., Lieslehto, J., Korhonen, V., Rytty, R., Mattila, H., Huotari, N., Raitamaa, L., Helakari, H., Elseoud, A. A., Krüger, J., LeVan, P., Tervonen, O., Hennig, J., Remes, A. M., Nedergaard, M., & Kiviniemi, V. (2020). The variability of functional MRI brain signal increases in Alzheimer’s disease at cardiorespiratory frequencies. Scientific Reports 2020 10:1, 10(1), 1–11. 10.1038/s41598-020-77984-1

Uludağ, K., Müller-Bierl, B., & Uğurbil, K. (2009). An integrative model for neuronal activity-induced signal changes for gradient and spin echo functional imaging. NeuroImage, 48(1), 150–165. 10.1016/J.NEUROIMAGE.2009.05.051

Urner, T. M., Cowdrick, K. R., Brothers, R. O., Boodooram, T., Zhao, H., Goyal, V., Sathialingam, E., Quadri, A., Turrentine, K., Akbar, M. M., Triplett, S. E., Bai, S., Buckley, E. M., Buckley, E. M., & Buckley, E. M. (2023). Normative cerebral microvascular blood flow waveform morphology assessed with diffuse correlation spectroscopy. Biomedical Optics Express, Vol. 14, Issue 7, Pp. 3635-3653, 14(7), 3635–3653. 10.1364/BOE.489760

Valsamis, J. J., Luciw, N. J., Haq, N., Atwi, S., Duchesne, S., Cameron, W., & MacIntosh, B. J. (2023). An imaging-based method of mapping multi-echo BOLD intracranial pulsatility. Magnetic Resonance in Medicine, 90(1), 343–352. 10.1002/MRM.29639

Van Bel, F., Van de Bor, M., Baan, J., Ruys, J. H., & Stijnen, T. (1988). The influence of abnormal blood gases on cerebral blood flow velocity in the preterm newborn. Neuropediatrics, 19(1), 27–32. 10.1055/S-2008-1052397/BIB

Van De Haar, H. J., Burgmans, S., Jansen, J. F. A., Van Osch, M. J. P., Van Buchem, M. A., Muller, M., Hofman, P. A. M., Verhey, F. R. J., & Backes, W. H. (2016). Blood-brain barrier leakage in patients with early Alzheimer disease. Radiology, 281(2), 527–535. 10.1148/radiol.2016152244

van den Kerkhof, M., Jansen, J. F. A., van Oostenbrugge, R. J., & Backes, W. H. (2023). 1D versus 3D blood flow velocity and pulsatility measurements of lenticulostriate arteries at 7T MRI. Magnetic Resonance Imaging, 96, 144–150. 10.1016/J.MRI.2022.12.005

van Hespen, K. M., Kuijf, H. J., Hendrikse, J., Luijten, P. R., & Zwanenburg, J. J. M. (2022). Blood Flow Velocity Pulsatility and Arterial Diameter Pulsatility Measurements of the Intracranial Arteries Using 4D PC-MRI. Neuroinformatics, 20(2), 317–326. 10.1007/S12021-021-09526-7/FIGURES/4

Van Hulst, E., Báez-Yáñez, M. G., Adams, A. L., Biessels, G. J., & Zwanenburg, J. J. M. (2024). The heartbeat induces local volumetric compression in the healthy human brain: a 7 T magnetic resonance imaging study on brain tissue pulsations. Interface Focus, 14(6). 10.1098/RSFS.2024.0032

van Tuijl, R. J., Ruigrok, Y. M., Velthuis, B. K., van der Schaaf, I. C., Rinkel, G. J. E., & Zwanenburg, J. J. M. (2020). Velocity pulsatility and arterial distensibility along the internal carotid artery. Journal of the American Heart Association, 9(16), 16883. 10.1161/JAHA.120.016883/SUPPL_FILE/JAH35395-SUP-0001-TABLES1.PDF

Viessmann, O., Möller, H. E., & Jezzard, P. (2017). Cardiac cycle-induced EPI time series fluctuations in the brain: Their temporal shifts, inflow effects and T2* fluctuations. NeuroImage, 162, 93–105. 10.1016/J.NEUROIMAGE.2017.08.061

Viessmann, O., Möller, H. E., & Jezzard, P. (2019). Dual regression physiological modeling of resting-state EPI power spectra: Effects of healthy aging. NeuroImage, 187, 68–76. 10.1016/J.NEUROIMAGE.2018.01.011

Vikner, T., Nyberg, L., Holmgren, M., Malm, J., Eklund, A., & Wåhlin, A. (2020). Characterizing pulsatility in distal cerebral arteries using 4D flow MRI. Journal of Cerebral Blood Flow and Metabolism, 40(12), 2429–2440. 10.1177/0271678X19886667

Wagshul, M. E., Eide, P. K., & Madsen, J. R. (2011a). The pulsating brain: A review of experimental and clinical studies of intracranial pulsatility. Fluids and Barriers of the CNS, 8(1), 1–23. 10.1186/2045-8118-8-5/FIGURES/12

Wagshul, M. E., Eide, P. K., & Madsen, J. R. (2011b). The pulsating brain: A review of experimental and clinical studies of intracranial pulsatility. Fluids and Barriers of the CNS, 8(1), 1–23. 10.1186/2045-8118-8-5/FIGURES/12

Wang, Q., Pan, M., Kreiss, L., Samaei, S., Carp, S. A., Johansson, J. D., Zhang, Y., Wu, M., Horstmeyer, R., Diop, M., & Li, D. D. U. (2024). A comprehensive overview of diffuse correlation spectroscopy: Theoretical framework, recent advances in hardware, analysis, and applications. NeuroImage, 298, 120793. 10.1016/J.NEUROIMAGE.2024.120793

Zarrinkoob, L., Ambarki, K., Wahlin, A., Birgander, R., Carlberg, B., Eklund, A., & Malm, J. (2016). Aging alters the dampening of pulsatile blood flow in cerebral arteries. Journal of Cerebral Blood Flow and Metabolism, 36(9), 1519–1527. 10.1177/0271678X16629486/SUPPL_FILE/SUPPLEMENTARY_MATERIAL.DOC

Zhao, F., Wang, P., Hendrich, K., Ugurbil, K., & Kim, S. G. (2006). Cortical layer-dependent BOLD and CBV responses measured by spin-echo and gradient-echo fMRI: Insights into hemodynamic regulation. NeuroImage, 30(4), 1149–1160. 10.1016/J.NEUROIMAGE.2005.11.013

Zhou, W., Kholiqov, O., Zhu, J., Zhao, M., Zimmermann, L. L., Martin, R. M., Lyeth, B. G., & Srinivasan, V. J. (2021). Functional interferometric diffusing wave spectroscopy of the human brain. Science Advances, 7(20), 150–162. 10.1126/SCIADV.ABE0150/SUPPL_FILE/ABE0150_SM.PDF

