## Supplementary Material for "Cerebrovascular pulsatility differs across vascular compartments and is altered by hypercapnic stimuli: a BOLD fMRI study"

### S1: CVR, CBV across cortical depth

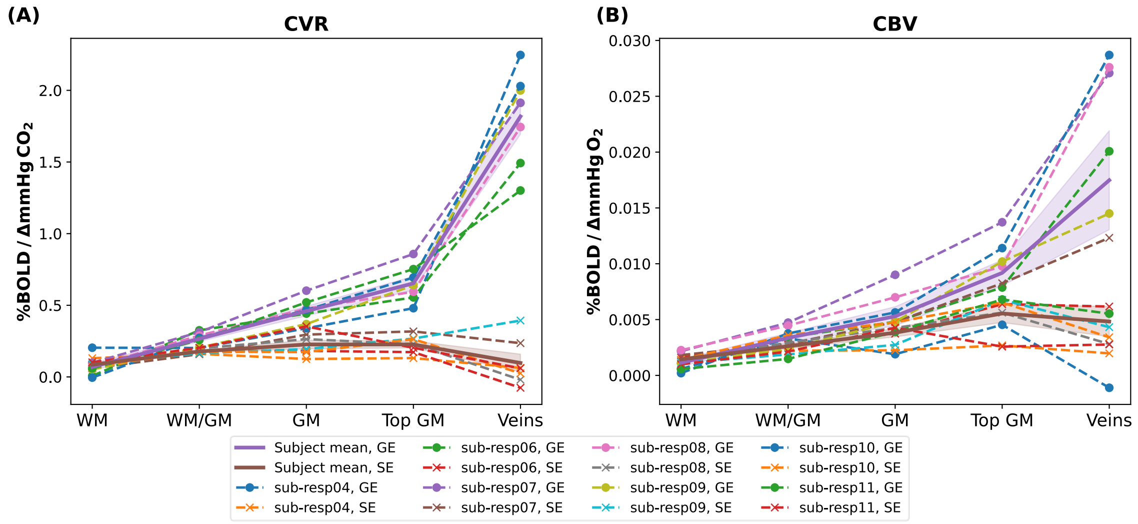


S1: $\mathrm{CVR}^{GE/SE}$ (A) and $\mathrm{CBV}v_{O_2}{}^{GE/SE}$ (B) across cortical depth. The quantities are plotted as a subject average (thick lines, shaded are represents standard error of mean) for Gradient-echo and Spin-echo.

### S2: PI across cortical depth (ROI average first)

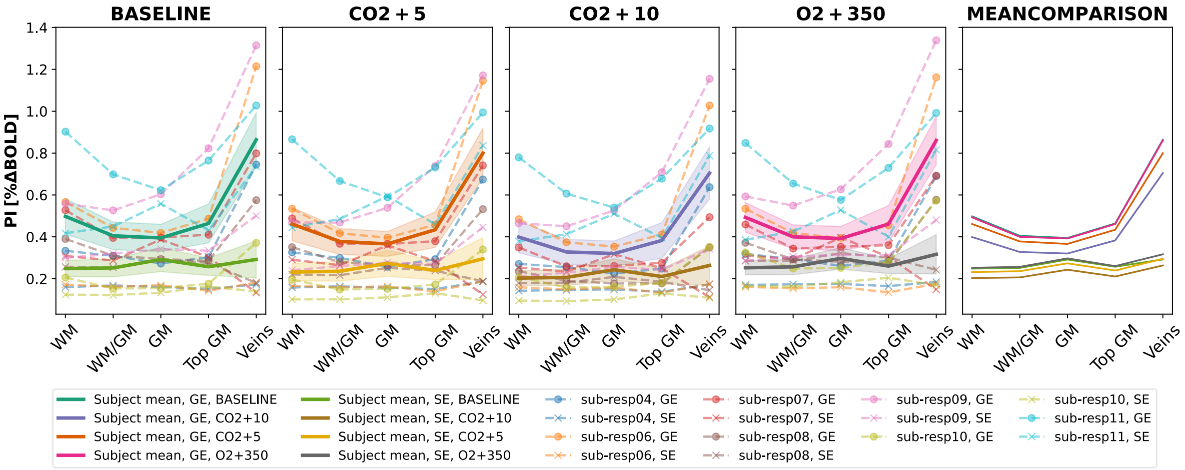


S2: Pulsatility Index ($PI$) across cortical depth. The pulsatility index (top row) is plotted across cortical depth at baseline, +5mmHg $PetCO_2$, +10mmHg $PetCO_2$, and +350mmHg $PetO_2$ (left to right). Mean quantities are computed per subject by aligning individual voxels gated time series temporally, computing a mean time series per ROI, and computing $PI$ based on the ROI-average gated time-series. Mean quantities are plotted for the individual subjects (dashed lines) and the subject average (thick lines, shaded area represents standard error of mean) for Gradient-echo and Spin-echo.

#### S3: Pulse Amplitude vs Mean Signal

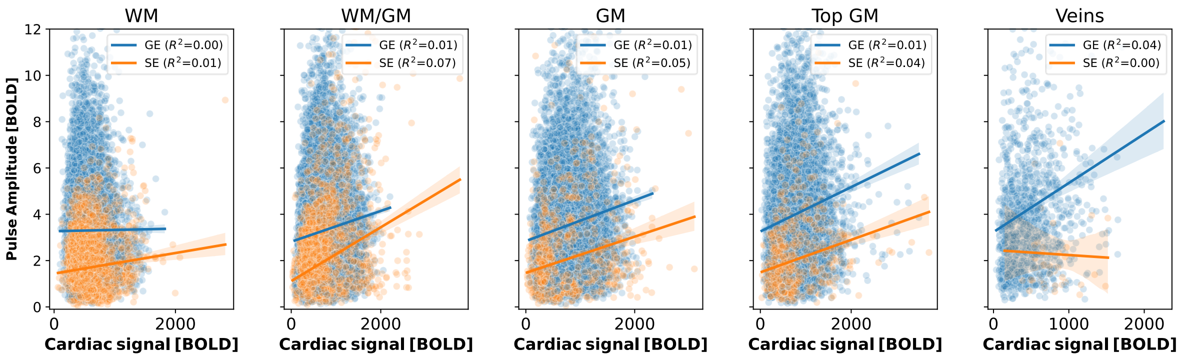


S3: A scatter plot and the corresponding linear regression (think line: mean signal, shaded area: standard error of the mean) between the Pulse Amplitude ($PA$) and mean cardiac signal is plotted for all ROIs (left to right: WM, deep GM, middle GM, top GM, and larger/pial veins).

#### S4: Cardiac Beats Per Minute

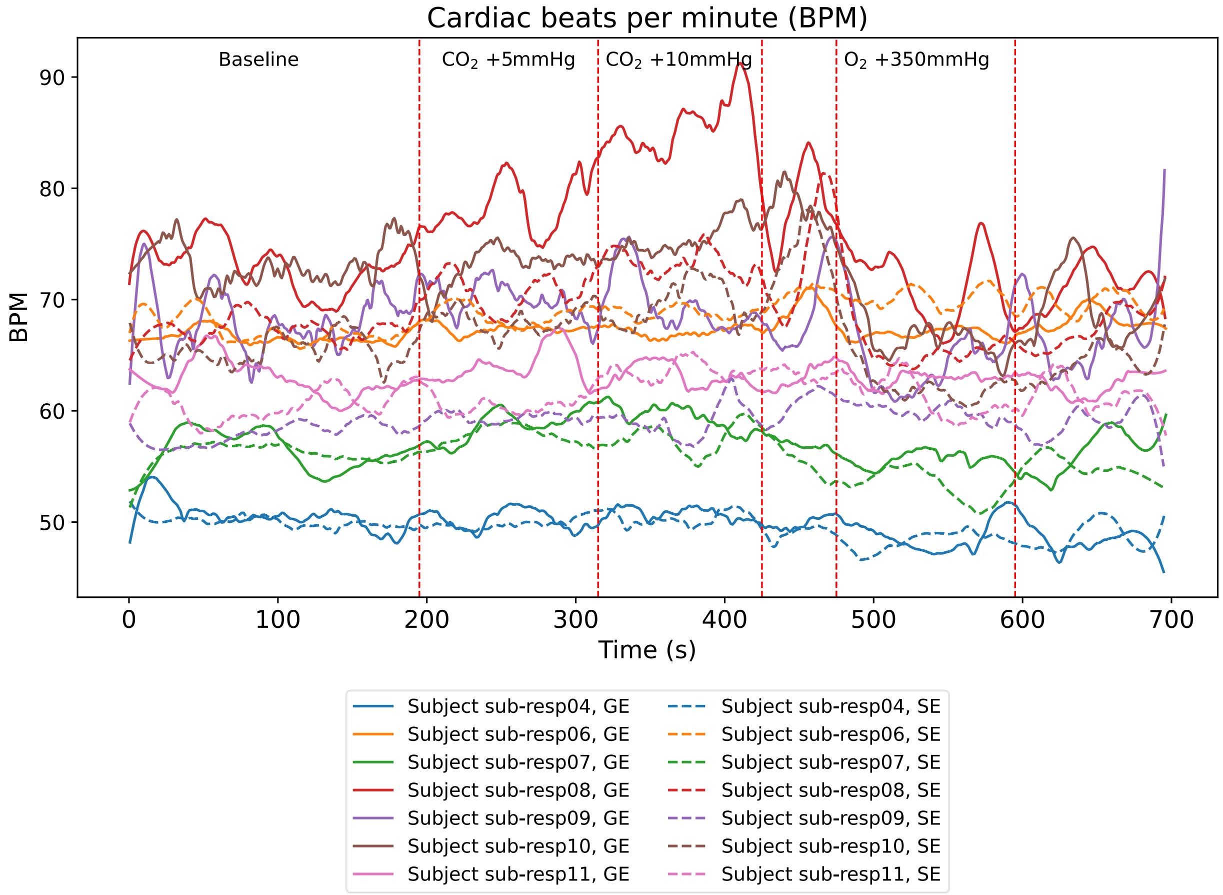


| Scan sequence | Subject ID | Baseline | CO2+5 | CO2+10 | O2+350 |
|---|---|---|---|---|---|
| GE | Subject 04 | 51 | 50 | 51 | 49 |
| GE | Subject 06 | 67 | 67 | 67 | 67 |
| GE | Subject 07 | 56 | 59 | 59 | 56 |
| GE | Subject 08 | 73 | 79 | 85 | 71 |
| GE | Subject 09 | 68 | 70 | 69 | 65 |
| GE | Subject 10 | 73 | 73 | 76 | 67 |
| GE | Subject 11 | 63 | 64 | 63 | 63 |
| SE | Subject 04 | 50 | 50 | 50 | 48 |
| SE | Subject 06 | 67 | 69 | 69 | 70 |
| SE | Subject 07 | 56 | 58 | 58 | 53 |
| SE | Subject 08 | 67 | 71 | 73 | 66 |
| SE | Subject 09 | 58 | 59 | 59 | 60 |
| SE | Subject 10 | 66 | 68 | 70 | 62 |
| SE | Subject 11 | 61 | 61 | 64 | 61 |

S4: Plot of the cardiac beats per minute (BPM) during the BOLD fMRI acquisition for GE (thick line) and SE (dashed line) for all subjects. In the table below, the mean BPM are reported for each subject during the baseline period, the two stages of hypercapnia (+5mmHg $PetCO_2$, +10mmHg $PetCO_2$), and the hyperoxia stage (+350mmHg $PetO_2$). With the exception of subject 08, no noticeable differences in the BPM's become apparent between the different breathing periods.

#### S5: Linear Mixed Model Fits

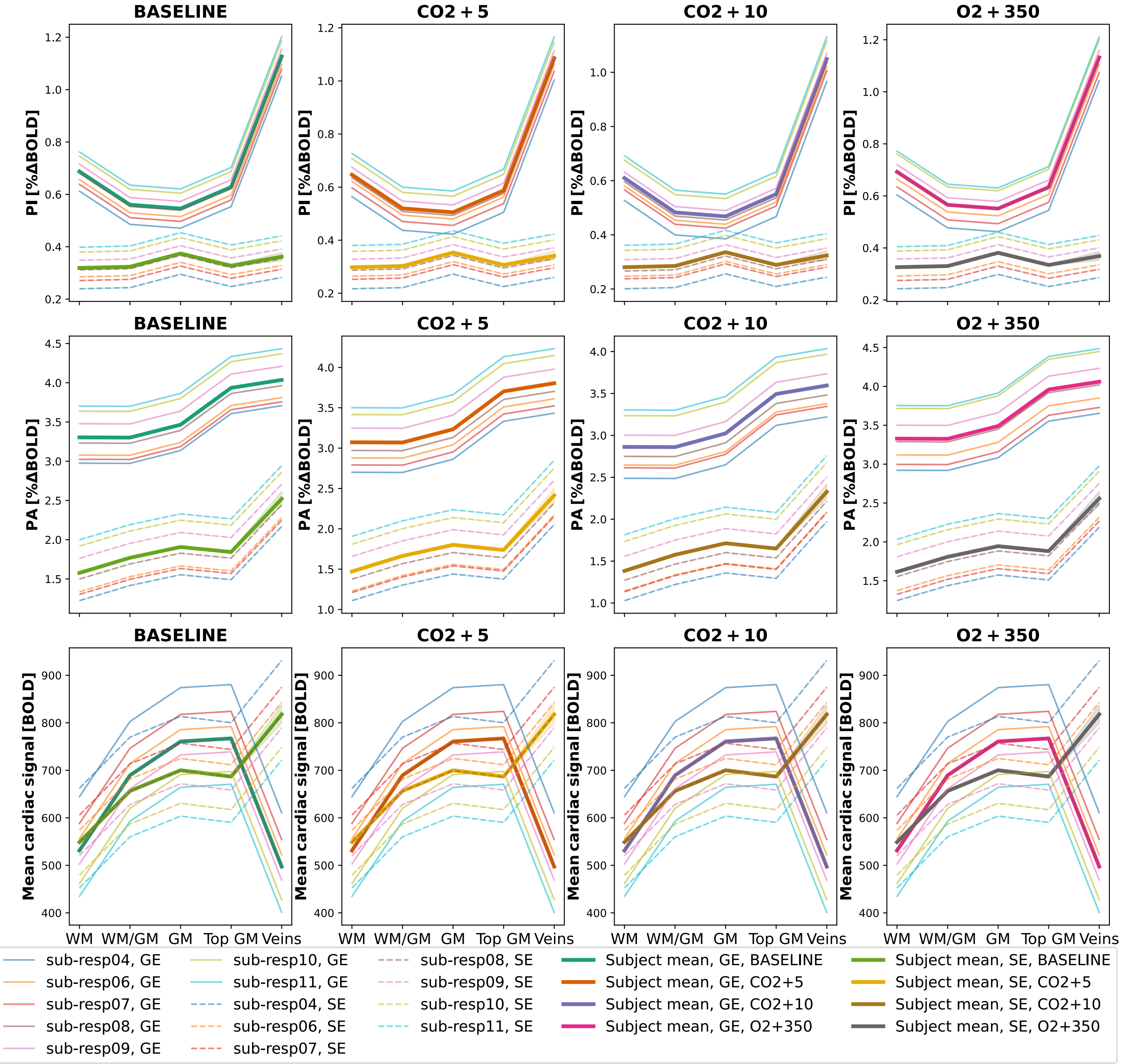

S5: Linear mixed model fits for the Pulsatility Index ($PI$), Pulse Amplitude ($PA$), and mean cardiac signal across cortical depth. The pulsatility index (top row, A), pulse amplitude (middle row, B), and cardiac BOLD signal (bottom row, C) are plotted across cortical depth at baseline, +5mmHg $PetCO_2$, +10mmHg $PetCO_2$, and +350mmHg $PetO_2$ (left to right). Mean quantities are plotted for the individual subjects (thin lines) and the subject average (thick lines) for Gradient-echo and Spin-echo.

#### S6: Statistical analysis of PI across cortical depth

##### Linear Mixed Model coefficients

| Variable | Estimate | Std. Error | df | t value | Pr(>\|t\|) |
|---|---|---|---|---|---|
| **(Intercept)** | 0.9432 | 1.6783 | 4.0006 | 0.5620 | 0.6041 |
| **scanTypeSE** | -0.3701 | 0.0453 | 6.0807 | -8.1656 | 0.0002 |
| **layerLoclayer_1** | -0.1269 | 0.0021 | 308198.0013 | -60.3384 | 0.0000 |
| **layerLoclayer_2** | -0.1416 | 0.0024 | 308198.0013 | -59.2122 | 0.0000 |
| **layerLoclayer_3** | -0.0594 | 0.0028 | 308198.0013 | -21.5770 | 0.0000 |
| **layerLoclayer_4** | 0.4392 | 0.0062 | 308198.0013 | 71.1962 | 0.0000 |
| **CO2_between** | -0.0150 | 0.0265 | 3.9943 | -0.5668 | 0.6012 |
| **O2_between** | 0.0035 | 0.0188 | 3.9899 | 0.1846 | 0.8625 |
| **CO2_ws** | -0.0089 | 0.0003 | 308201.2905 | -34.4667 | 0.0000 |
| **O2_ws** | 0.0000 | 0.0000 | 308201.0754 | -2.2149 | 0.0268 |
| **scanTypeSE:layerLoclayer_1** | 0.1314 | 0.0041 | 308198.0013 | 32.2950 | 0.0000 |
| **scanTypeSE:layerLoclayer_2** | 0.1969 | 0.0055 | 308198.0013 | 36.0419 | 0.0000 |
| **scanTypeSE:layerLoclayer_3** | 0.0679 | 0.0061 | 308198.0013 | 11.1287 | 0.0000 |
| **scanTypeSE:layerLoclayer_4** | -0.3962 | 0.0315 | 308198.0013 | -12.5696 | 0.0000 |
| **scanTypeSE:CO2_ws** | 0.0043 | 0.0006 | 308167.4464 | 7.7156 | 0.0000 |
| **scanTypeSE:O2_ws** | 0.0000 | 0.0000 | 308191.4413 | 1.1621 | 0.2452 |

##### LRT and PBtest Summary

| Iteration | Tested_Variable | LRT_statistic | LRT_df | LRT_p_value | PB_statistic | PB_p_value |
|---|---|---|---|---|---|---|
| **1** | scanType | 486.9246 | 9.0000 | 0.0000 | 486.9246 | 0.0001 |
| **2** | layerLoc | 347.8234 | 8.0000 | 0.0000 | 347.8234 | 0.0001 |
| **3** | scanType:layerLoc | 217.0551 | 4.0000 | 0.0000 | 217.0551 | 0.0001 |
| **4** | CO2_between | 0.7477 | 1.0000 | 0.3872 | 0.7477 | 0.5933 |
| **5** | CO2_ws | 19.3660 | 2.0000 | 0.0001 | 19.3660 | 0.0001 |
| **6** | O2_between | 0.2193 | 1.0000 | 0.6396 | 0.2193 | 0.7772 |
| **7** | O2_ws | 0.0256 | 2.0000 | 0.9873 | 0.0256 | 0.9872 |
| **8** | CO2_ws:scanType | 2.3360 | 1.0000 | 0.1264 | 2.3360 | 0.1362 |
| **9** | O2_ws:scanType | 0.0081 | 1.0000 | 0.9283 | 0.0081 | 0.9339 |

##### Post-hoc Test: Laminae comparison

| Contrast | scanType | estimate | SE | df | t.ratio | p.value |
|---|---|---|---|---|---|---|
| **layer_0 - layer_1** | GE | 0.1269 | 0.0261 | 254.0000 | 4.8707 | 0.0000 |
| **layer_0 - layer_2** | GE | 0.1416 | 0.0261 | 254.0000 | 5.4324 | 0.0000 |
| **layer_0 - layer_3** | GE | 0.0594 | 0.0261 | 254.0000 | 2.2782 | 0.1554 |
| **layer_0 - layer_4** | GE | -0.4392 | 0.0261 | 254.0000 | -16.8547 | 0.0000 |
| **layer_1 - layer_2** | GE | 0.0146 | 0.0261 | 254.0000 | 0.5617 | 0.9804 |
| **layer_1 - layer_3** | GE | -0.0676 | 0.0261 | 254.0000 | -2.5925 | 0.0748 |

| | | | | | | |
|---|---|---|---|---|---|---|
| **layer_1 - layer_4** | GE | -0.5662 | 0.0261 | 254.0000 | -21.7254 | 0.0000 |
| **layer_2 - layer_3** | GE | -0.0822 | 0.0261 | 254.0000 | -3.1542 | 0.0154 |
| **layer_2 - layer_4** | GE | -0.5808 | 0.0261 | 254.0000 | -22.2871 | 0.0000 |
| **layer_3 - layer_4** | GE | -0.4986 | 0.0261 | 254.0000 | -19.1329 | 0.0000 |
| **layer_0 - layer_1** | SE | -0.0045 | 0.0261 | 254.0000 | -0.1730 | 0.9998 |
| **layer_0 - layer_2** | SE | -0.0553 | 0.0261 | 254.0000 | -2.1239 | 0.2131 |
| **layer_0 - layer_3** | SE | -0.0085 | 0.0261 | 254.0000 | -0.3268 | 0.9975 |

#### Post-hoc Test: Gas Level comparison

| contrast | scanType | estimate | SE | df | t.ratio | p.value |
|---|---|---|---|---|---|---|
| **BASELINE - (CO2+10)** | GE | 0.0920 | 0.0233 | 252.0000 | 3.9456 | 0.0006 |
| **BASELINE - (CO2+5)** | GE | 0.0139 | 0.0233 | 252.0000 | 0.5964 | 0.9331 |
| **BASELINE - (O2+350)** | GE | -0.0187 | 0.0233 | 252.0000 | -0.8018 | 0.8535 |
| **(CO2+10) - (CO2+5)** | GE | -0.0781 | 0.0233 | 252.0000 | -3.3492 | 0.0051 |
| **(CO2+10) - (O2+350)** | GE | -0.1106 | 0.0233 | 252.0000 | -4.7474 | 0.0000 |
| **(CO2+5) - (O2+350)** | GE | -0.0326 | 0.0233 | 252.0000 | -1.3982 | 0.5016 |
| **BASELINE - (CO2+10)** | SE | 0.0397 | 0.0233 | 252.0000 | 1.7038 | 0.3238 |
| **BASELINE - (CO2+5)** | SE | 0.0129 | 0.0233 | 252.0000 | 0.5553 | 0.9450 |
| **BASELINE - (O2+350)** | SE | -0.0080 | 0.0233 | 252.0000 | -0.3439 | 0.9860 |
| **(CO2+10) - (CO2+5)** | SE | -0.0268 | 0.0233 | 252.0000 | -1.1485 | 0.6598 |
| **(CO2+10) - (O2+350)** | SE | -0.0477 | 0.0233 | 252.0000 | -2.0477 | 0.1735 |
| **(CO2+5) - (O2+350)** | SE | -0.0210 | 0.0233 | 252.0000 | -0.8992 | 0.8052 |

### S7: Statistical Analysis of PA across cortical depth

#### Linear Mixed Model Coefficients

| Variable | Estimate | Std. Error | df | t value | Pr(>\|t\|) |
|---|---|---|---|---|---|
| **(Intercept)** | 6.0662 | 9.4397 | 4.0149 | 0.6426 | 0.5553 |
| **scanTypeSE** | -1.7368 | 0.3564 | 6.0123 | -4.8736 | 0.0028 |
| **layerLoclayer_1** | -0.0025 | 0.0091 | 308198.0009 | -0.2731 | 0.7848 |
| **layerLoclayer_2** | 0.1614 | 0.0103 | 308198.0008 | 15.6449 | 0.0000 |
| **layerLoclayer_3** | 0.6314 | 0.0119 | 308198.0008 | 53.1777 | 0.0000 |
| **layerLoclayer_4** | 0.7321 | 0.0266 | 308198.0008 | 27.5019 | 0.0000 |
| **CO2_between** | -0.0607 | 0.1491 | 3.9981 | -0.4070 | 0.7048 |
| **O2_between** | 0.0342 | 0.1059 | 3.9969 | 0.3225 | 0.7632 |
| **CO2_ws** | -0.0504 | 0.0011 | 308199.0277 | -45.4026 | 0.0000 |
| **O2_ws** | -0.0001 | 0.0000 | 308198.9324 | -3.5190 | 0.0004 |
| **scanTypeSE:layerLoclayer_1** | 0.1950 | 0.0176 | 308198.0008 | 11.1041 | 0.0000 |
| **scanTypeSE:layerLoclayer_2** | 0.1686 | 0.0236 | 308198.0008 | 7.1520 | 0.0000 |
| **scanTypeSE:layerLoclayer_3** | -0.3650 | 0.0263 | 308198.0008 | -13.8658 | 0.0000 |
| **scanTypeSE:layerLoclayer_4** | 0.2114 | 0.1360 | 308198.0008 | 1.5544 | 0.1201 |
| **scanTypeSE:CO2_ws** | 0.0275 | 0.0024 | 308205.3482 | 11.3681 | 0.0000 |

| | | | | | |
|---|---|---|---|---|---|
| **scanTypeSE:O2_ws** | 0.0001 | 0.0001 | 308204.6123 | 1.9435 | 0.0520 |

#### LRT and PBtest Summary

| Iteration | Tested_Variable | LRT_statistic | LRT_df | LRT_p_value | PB_statistic | PB_p_value |
|---|---|---|---|---|---|---|
| **1** | scanType | 363.5574 | 9.0000 | 0.0000 | 363.5574 | 0.0001 |
| **2** | layerLoc | 81.1390 | 8.0000 | 0.0000 | 81.1390 | 0.0001 |
| **3** | scanType:layerLoc | 10.9014 | 4.0000 | 0.0277 | 10.9014 | 0.0298 |
| **4** | CO2_between | 0.5200 | 1.0000 | 0.4708 | 0.5200 | 0.6624 |
| **5** | CO2_ws | 15.8629 | 2.0000 | 0.0004 | 15.8629 | 0.0007 |
| **6** | O2_between | 0.1228 | 1.0000 | 0.7260 | 0.1228 | 0.8328 |
| **7** | O2_ws | 0.0531 | 2.0000 | 0.9738 | 0.0531 | 0.9754 |
| **8** | CO2_ws:scanType | 1.5311 | 1.0000 | 0.2159 | 1.5311 | 0.2218 |
| **9** | O2_ws:scanType | 0.0156 | 1.0000 | 0.9006 | 0.0156 | 0.9055 |

### S8: Statistical Analysis of mean cardiac BOLD signal across cortical depth

#### Linear Mixed Model coefficients

| Variable | Estimate | Std. Error | df | t value | Pr(>\|t\|) |
|---|---|---|---|---|---|
| **(Intercept)** | 552.5427 | 511.5217 | 4.0123 | 1.0802 | 0.3407 |
| **scanTypeSE** | 17.6468 | 37.8565 | 6.0137 | 0.4661 | 0.6575 |
| **layerLoclayer_1** | 158.2937 | 1.2998 | 308199.9999 | 121.7847 | 0.0000 |
| **layerLoclayer_2** | 229.5561 | 1.4772 | 308199.9999 | 155.3949 | 0.0000 |
| **layerLoclayer_3** | 236.0265 | 1.7001 | 308199.9999 | 138.8321 | 0.0000 |
| **layerLoclayer_4** | -33.8545 | 3.8119 | 308199.9999 | -8.8813 | 0.0000 |
| **CO2_between** | 23.6010 | 8.0803 | 3.9997 | 2.9208 | 0.0432 |
| **CO2_ws** | -0.0012 | 0.1411 | 308204.7149 | -0.0086 | 0.9931 |
| **O2_between** | -0.5471 | 5.7396 | 3.9992 | -0.0953 | 0.9287 |
| **O2_ws** | 0.0000 | 0.0038 | 308203.9957 | 0.0047 | 0.9962 |
| **scanTypeSE:layerLoclayer_1** | -50.6079 | 2.5147 | 308199.9999 | -20.1246 | 0.0000 |
| **scanTypeSE:layerLoclayer_2** | -78.2227 | 3.3758 | 308199.9999 | -23.1717 | 0.0000 |
| **scanTypeSE:layerLoclayer_3** | -97.9923 | 3.7690 | 308199.9999 | -25.9995 | 0.0000 |
| **scanTypeSE:layerLoclayer_4** | 302.9522 | 19.4735 | 308199.9999 | 15.5572 | 0.0000 |

#### LRT and PBtest Summary

| Iteration | Tested_Variable | LRT_statistic | LRT_df | LRT_p_value | PB_statistic | PB_p_value |
|---|---|---|---|---|---|---|
| **1** | scanType | 246.8061 | 9.0000 | 0.0000 | 246.8061 | 0.0001 |
| **2** | layerLoc | 324.5577 | 8.0000 | 0.0000 | 324.5577 | 0.0001 |
| **3** | scanType:layerLoc | 218.3265 | 4.0000 | 0.0000 | 218.3265 | 0.0001 |
| **4** | CO2_between | 6.1943 | 1.0000 | 0.0128 | 6.1943 | 0.1183 |

| | | | | | | |
|---|---|---|---|---|---|---|
| **5** | CO2_ws | 0.0112 | 2.0000 | 0.9944 | 0.0112 | 0.9940 |
| **6** | O2_between | 0.9302 | 1.0000 | 0.3348 | 0.9302 | 0.5417 |
| **7** | O2_ws | 0.0113 | 2.0000 | 0.9944 | 0.0113 | 0.9944 |
| **8** | CO2_ws:scanType | 0.0056 | 1.0000 | 0.9402 | 0.0056 | 0.9432 |
| **9** | O2_ws:scanType | 0.0065 | 1.0000 | 0.9355 | 0.0065 | 0.9389 |

### S9: Statistical Analysis of $\Delta PI_{CO2}$ across cortical depth

#### Linear Mixed Model coefficients

| Variable | Estimate | Std. Error | df | t value | Pr(>\|t\|) |
|---|---|---|---|---|---|
| **(Intercept)** | -0.0098 | 0.0024 | 6.0103 | -4.0672 | 0.0066 |
| **scanTypeSE** | 0.0054 | 0.0017 | 6.0586 | 3.1517 | 0.0195 |
| **layerLoclayer_1** | 0.0018 | 0.0001 | 77034.0001 | 15.8824 | 0.0000 |
| **layerLoclayer_2** | 0.0019 | 0.0001 | 77034.0001 | 14.7945 | 0.0000 |
| **layerLoclayer_3** | 0.0013 | 0.0001 | 77034.0001 | 9.0259 | 0.0000 |
| **layerLoclayer_4** | -0.0065 | 0.0003 | 77034.0001 | -19.5615 | 0.0000 |
| **scanTypeSE:layerLoclayer_1** | -0.0017 | 0.0002 | 77034.0001 | -7.8507 | 0.0000 |
| **scanTypeSE:layerLoclayer_2** | -0.0024 | 0.0003 | 77034.0001 | -8.0987 | 0.0000 |
| **scanTypeSE:layerLoclayer_3** | -0.0014 | 0.0003 | 77034.0001 | -4.2408 | 0.0000 |
| **scanTypeSE:layerLoclayer_4** | 0.0058 | 0.0017 | 77034.0001 | 3.3920 | 0.0007 |

#### LRT and PBtest Summary

| Iteration | Tested_Variable | LRT_statistic | LRT_df | LRT_p_value | PB_statistic | PB_p_value |
|---|---|---|---|---|---|---|
| **1** | scanType | 90.8717 | 7.0000 | 0.0000 | 90.8717 | 0.0001 |
| **2** | layerLoc | 50.7399 | 8.0000 | 0.0000 | 50.7399 | 0.0001 |
| **3** | scanType:layerLoc | 27.4398 | 4.0000 | 0.0000 | 27.4398 | 0.0002 |

#### Post-hoc Test: Laminae comparison

| contrast | scanType | estimate | SE | df | t.ratio | p.value |
|---|---|---|---|---|---|---|
| **layer_0 - layer_1** | GE | -0.0018 | 0.0012 | 48.0000 | -1.4993 | 0.5680 |
| **layer_0 - layer_2** | GE | -0.0019 | 0.0012 | 48.0000 | -1.5873 | 0.5126 |
| **layer_0 - layer_3** | GE | -0.0013 | 0.0012 | 48.0000 | -1.1144 | 0.7981 |
| **layer_0 - layer_4** | GE | 0.0065 | 0.0012 | 48.0000 | 5.4155 | 0.0000 |
| **layer_1 - layer_2** | GE | -0.0001 | 0.0012 | 48.0000 | -0.0880 | 1.0000 |
| **layer_1 - layer_3** | GE | 0.0005 | 0.0012 | 48.0000 | 0.3848 | 0.9952 |
| **layer_1 - layer_4** | GE | 0.0083 | 0.0012 | 48.0000 | 6.9148 | 0.0000 |
| **layer_2 - layer_3** | GE | 0.0006 | 0.0012 | 48.0000 | 0.4728 | 0.9895 |
| **layer_2 - layer_4** | GE | 0.0084 | 0.0012 | 48.0000 | 7.0027 | 0.0000 |
| **layer_3 - layer_4** | GE | 0.0078 | 0.0012 | 48.0000 | 6.5299 | 0.0000 |
| **layer_0 - layer_1** | SE | -0.0001 | 0.0012 | 48.0000 | -0.0655 | 1.0000 |

| | | | | | | |
|---|---|---|---|---|---|---|
| **layer_0 - layer_2** | SE | 0.0005 | 0.0012 | 48.0000 | 0.3983 | 0.9945 |
| **layer_0 - layer_3** | SE | 0.0001 | 0.0012 | 48.0000 | 0.0464 | 1.0000 |
| **layer_0 - layer_4** | SE | 0.0007 | 0.0012 | 48.0000 | 0.6181 | 0.9715 |
| **layer_1 - layer_2** | SE | 0.0006 | 0.0012 | 48.0000 | 0.4638 | 0.9902 |
| **layer_1 - layer_3** | SE | 0.0001 | 0.0012 | 48.0000 | 0.1119 | 1.0000 |
| **layer_1 - layer_4** | SE | 0.0008 | 0.0012 | 48.0000 | 0.6836 | 0.9591 |
| **layer_2 - layer_3** | SE | -0.0004 | 0.0012 | 48.0000 | -0.3519 | 0.9966 |
| **layer_2 - layer_4** | SE | 0.0003 | 0.0012 | 48.0000 | 0.2198 | 0.9995 |
| **layer_3 - layer_4** | SE | 0.0007 | 0.0012 | 48.0000 | 0.5717 | 0.9786 |

### S10: Statistical Analysis of PI vs CVR

#### Linear Mixed Model coefficients

| Variable | Estimate | Std. Error | df | t value | Pr(>\|t\|) |
|---|---|---|---|---|---|
| **(Intercept)** | 0.6015 | 0.0618 | 6.0964 | 9.7392 | 0.0001 |
| **CVR** | 0.0250 | 0.0045 | 8.0064 | 5.5156 | 0.0006 |
| **scanTypeSE** | -0.3336 | 0.0077 | 55515.8373 | -43.1069 | 0.0000 |
| **layerLoclayer_1** | -0.1471 | 0.0073 | 55457.5874 | -20.2575 | 0.0000 |
| **layerLoclayer_2** | -0.2290 | 0.0080 | 55292.3985 | -28.7705 | 0.0000 |
| **layerLoclayer_3** | -0.1440 | 0.0086 | 54354.2475 | -16.7373 | 0.0000 |
| **layerLoclayer_4** | 0.0191 | 0.0200 | 55071.4162 | 0.9567 | 0.3387 |
| **CVR:scanTypeSE** | 0.0006 | 0.0013 | 55398.0935 | 0.4485 | 0.6538 |
| **CVR:layerLoclayer_1** | -0.0047 | 0.0018 | 54790.2290 | -2.5755 | 0.0100 |
| **CVR:layerLoclayer_2** | 0.0021 | 0.0018 | 52809.0475 | 1.1411 | 0.2538 |
| **CVR:layerLoclayer_3** | -0.0057 | 0.0018 | 51991.4407 | -3.1591 | 0.0016 |
| **CVR:layerLoclayer_4** | -0.0008 | 0.0018 | 55090.6931 | -0.4326 | 0.6653 |
| **scanTypeSE:layerLoclayer_1** | 0.1336 | 0.0100 | 55515.9829 | 13.3602 | 0.0000 |
| **scanTypeSE:layerLoclayer_2** | 0.2360 | 0.0133 | 55515.8902 | 17.7656 | 0.0000 |
| **scanTypeSE:layerLoclayer_3** | 0.1262 | 0.0143 | 55511.7530 | 8.8345 | 0.0000 |
| **scanTypeSE:layerLoclayer_4** | 0.0657 | 0.0906 | 55512.4902 | 0.7251 | 0.4684 |

#### LRT and PBtest Summary

| Iteration | Tested_Variable | LRT_statistic | LRT_df | LRT_p_value | PB_statistic | PB_p_value |
|---|---|---|---|---|---|---|
| **1** | scanType | 29.1196 | 6.0000 | 0.0001 | 29.1196 | 0.0009 |
| **2** | layerLoc | 23.0221 | 12.0000 | 0.0275 | 23.0221 | 0.0948 |
| **3** | CVR | 34.3749 | 8.0000 | 0.0000 | 34.3749 | 0.0006 |
| **4** | CVR:scanType | 0.9700 | 1.0000 | 0.3247 | 0.9700 | 0.4037 |
| **5** | CVR:layerLoc | 5.1482 | 4.0000 | 0.2724 | 5.1482 | 0.4282 |
| **6** | scanType:layerLoc | 3.6287 | 4.0000 | 0.4586 | 3.6287 | 0.6036 |

### S11: Statistical Analysis of PI vs CBV

#### Linear Mixed Model coefficients

| Variable | Estimate | Std. Error | df | t value | Pr(>\|t\|) |
|---|---|---|---|---|---|
| **(Intercept)** | 0.5582 | 0.0576 | 6.1506 | 9.6822 | 0.0001 |
| **CBV** | 0.0295 | 0.0053 | 6.9519 | 5.5519 | 0.0009 |
| **scanTypeSE** | -0.3196 | 0.0078 | 55513.3791 | -41.0010 | 0.0000 |
| **layerLoclayer_1** | -0.0975 | 0.0079 | 55485.4738 | -12.4146 | 0.0000 |
| **layerLoclayer_2** | -0.1750 | 0.0084 | 55445.9442 | -20.7872 | 0.0000 |
| **layerLoclayer_3** | -0.1403 | 0.0093 | 55287.4271 | -15.1488 | 0.0000 |
| **layerLoclayer_4** | 0.2130 | 0.0198 | 55459.8359 | 10.7570 | 0.0000 |
| **CBV:scanTypeSE** | -0.0040 | 0.0009 | 55514.0842 | -4.5961 | 0.0000 |
| **CBV:layerLoclayer_1** | -0.0123 | 0.0015 | 55399.6976 | -7.9268 | 0.0000 |
| **CBV:layerLoclayer_2** | -0.0067 | 0.0015 | 55210.2221 | -4.3648 | 0.0000 |
| **CBV:layerLoclayer_3** | -0.0101 | 0.0015 | 55167.3866 | -6.6285 | 0.0000 |
| **CBV:layerLoclayer_4** | -0.0129 | 0.0016 | 55283.5428 | -8.2256 | 0.0000 |
| **scanTypeSE:layerLoclayer_1** | 0.1185 | 0.0100 | 55514.7177 | 11.8738 | 0.0000 |
| **scanTypeSE:layerLoclayer_2** | 0.2054 | 0.0133 | 55514.6211 | 15.4311 | 0.0000 |
| **scanTypeSE:layerLoclayer_3** | 0.1191 | 0.0145 | 55515.5048 | 8.1925 | 0.0000 |
| **scanTypeSE:layerLoclayer_4** | -0.0906 | 0.0905 | 55510.5609 | -1.0007 | 0.3170 |

#### LRT and PBtest Summary

| Iteration | Tested_Variable | LRT_statistic | LRT_df | LRT_p_value | PB_statistic | PB_p_value |
|---|---|---|---|---|---|---|
| **1** | scanType | 36.77884 | 6 | 1.94E-06 | 36.77884 | 1.00E-04 |
| **2** | layerLoc | 27.44826 | 12 | 0.006657 | 27.44826 | 0.034897 |
| **3** | CBV | 39.96959 | 8 | 3.25E-06 | 39.96959 | 0.0002 |
| **4** | CBV:scanType | 6.710524 | 1 | 0.009585 | 6.710524 | 0.025685 |
| **5** | CBV:layerLoc | 1.369926 | 4 | 0.849405 | 1.369926 | 0.906635 |
| **6** | scanType:layerLoc | 9.825056 | 4 | 0.04348 | 9.825056 | 0.11634 |

### S12: Statistical Analysis of $\Delta PI_{CO2}$ vs CVR

#### Linear Mixed Model coefficients

| Variable | Estimate | Std. Error | df | t value | Pr(>\|t\|) |
|---|---|---|---|---|---|
| **(Intercept)** | -0.0094 | 0.0016 | 6.1132 | -6.0298 | 0.0009 |
| **CVR** | -0.0003 | 0.0001 | 8.0106 | -2.2780 | 0.0522 |
| **scanTypeSE** | 0.0056 | 0.0002 | 55459.2296 | 26.4844 | 0.0000 |
| **layerLoclayer_1** | 0.0022 | 0.0002 | 55462.7620 | 11.3763 | 0.0000 |
| **layerLoclayer_2** | 0.0041 | 0.0002 | 55464.3135 | 19.0626 | 0.0000 |
| **layerLoclayer_3** | 0.0036 | 0.0002 | 55463.6515 | 15.1731 | 0.0000 |
| **layerLoclayer_4** | 0.0014 | 0.0006 | 55465.1311 | 2.5431 | 0.0110 |

| | | | | | |
|---|---|---|---|---|---|
| **CVR:scanTypeSE** | 0.0000 | 0.0000 | 55463.5626 | -0.5369 | 0.5913 |
| **CVR:layerLoclayer_1** | 0.0001 | 0.0000 | 55462.2040 | 2.2529 | 0.0243 |
| **CVR:layerLoclayer_2** | -0.0002 | 0.0000 | 55442.4356 | -3.1002 | 0.0019 |
| **CVR:layerLoclayer_3** | 0.0000 | 0.0000 | 55435.2245 | -0.8565 | 0.3917 |
| **CVR:layerLoclayer_4** | -0.0001 | 0.0000 | 55464.4255 | -2.5185 | 0.0118 |
| **scanTypeSE:layerLoclayer_1** | -0.0021 | 0.0003 | 55459.1603 | -7.8537 | 0.0000 |
| **scanTypeSE:layerLoclayer_2** | -0.0035 | 0.0004 | 55459.0691 | -9.7661 | 0.0000 |
| **scanTypeSE:layerLoclayer_3** | -0.0028 | 0.0004 | 55460.1590 | -7.2734 | 0.0000 |
| **scanTypeSE:layerLoclayer_4** | -0.0022 | 0.0023 | 55458.5882 | -0.9825 | 0.3259 |

#### LRT and PBtest Summary

| Iteration | Tested_Variable | LRT_statistic | LRT_df | LRT_p_value | PB_statistic | PB_p_value |
|---|---|---|---|---|---|---|
| **1** | scanType | 19.4582 | 6.0000 | 0.0035 | 19.4582 | 0.0199 |
| **2** | layerLoc | 15.3160 | 12.0000 | 0.2246 | 15.3160 | 0.3978 |
| **3** | CVR | 38.7664 | 7.0000 | 0.0000 | 38.7664 | 0.0002 |
| **4** | CVR:scanType | 0.2429 | 1.0000 | 0.6221 | 0.2429 | 0.6732 |
| **5** | CVR:layerLoc | 3.6285 | 4.0000 | 0.4586 | 3.6285 | 0.5970 |
| **6** | scanType:layerLoc | 2.3424 | 4.0000 | 0.6731 | 2.3424 | 0.7680 |

### S13: Statistical Analysis of $\Delta PI_{CO2}$ vs CBV

#### Linear Mixed Model coefficients

| Variable | Estimate | Std. Error | df | t value | Pr(>\|t\|) |
|---|---|---|---|---|---|
| **(Intercept)** | -0.0085 | 0.0016 | 6.1448 | -5.2496 | 0.0018 |
| **CBV** | -0.0004 | 0.0001 | 10.7389 | -5.1772 | 0.0003 |
| **scanTypeSE** | 0.0053 | 0.0002 | 55460.9671 | 25.0697 | 0.0000 |
| **layerLoclayer_1** | 0.0015 | 0.0002 | 55458.6976 | 6.8805 | 0.0000 |
| **layerLoclayer_2** | 0.0029 | 0.0002 | 55447.4820 | 12.5696 | 0.0000 |
| **layerLoclayer_3** | 0.0032 | 0.0003 | 55376.7172 | 12.6717 | 0.0000 |
| **layerLoclayer_4** | -0.0026 | 0.0005 | 55443.7684 | -4.7048 | 0.0000 |
| **CBV:scanTypeSE** | 0.0000 | 0.0000 | 55462.6589 | 1.3681 | 0.1713 |
| **CBV:layerLoclayer_1** | 0.0002 | 0.0000 | 55427.9943 | 5.7466 | 0.0000 |
| **CBV:layerLoclayer_2** | 0.0001 | 0.0000 | 55374.5544 | 2.3725 | 0.0177 |
| **CBV:layerLoclayer_3** | 0.0001 | 0.0000 | 55358.7516 | 2.9213 | 0.0035 |
| **CBV:layerLoclayer_4** | 0.0002 | 0.0000 | 55432.3584 | 4.2601 | 0.0000 |
| **scanTypeSE:layerLoclayer_1** | -0.0020 | 0.0003 | 55461.7313 | -7.2372 | 0.0000 |
| **scanTypeSE:layerLoclayer_2** | -0.0029 | 0.0004 | 55461.4782 | -8.0661 | 0.0000 |
| **scanTypeSE:layerLoclayer_3** | -0.0024 | 0.0004 | 55463.0656 | -6.0395 | 0.0000 |
| **scanTypeSE:layerLoclayer_4** | 0.0009 | 0.0023 | 55458.3927 | 0.3844 | 0.7007 |

##### LRT and PBtest Summary

| Iteration | Tested_Variable | LRT_statistic | LRT_df | LRT_p_value | PB_statistic | PB_p_value |
|---|---|---|---|---|---|---|
| **1** | scanType | 16.9385 | 6.0000 | 0.0095 | 16.9385 | 0.0389 |
| **2** | layerLoc | 20.0724 | 12.0000 | 0.0657 | 20.0724 | 0.1683 |
| **3** | CBV | 40.1897 | 7.0000 | 0.0000 | 40.1897 | 0.0001 |
| **4** | CBV:scanType | 1.4846 | 1.0000 | 0.2231 | 1.4846 | 0.2917 |
| **5** | CBV:layerLoc | 6.5097 | 4.0000 | 0.1642 | 6.5097 | 0.2941 |
| **6** | scanType:layerLoc | 2.8506 | 4.0000 | 0.5831 | 2.8506 | 0.7047 |

#### S14: Statistical Analysis of $\Delta PI_{O2}$ across cortical depth

##### Linear Mixed Model coefficients

| Variable | Estimate | Std. Error | df | t value | Pr(>\|t\|) |
|---|---|---|---|---|---|
| **(Intercept)** | 0.0001 | 0.0001 | 6.0101 | 0.8957 | 0.4048 |
| **scanTypeSE** | 0.0000 | 0.0000 | 6.0523 | -0.7288 | 0.4933 |
| **layerLoclayer_1** | 0.0000 | 0.0000 | 77034.0001 | -4.3623 | 0.0000 |
| **layerLoclayer_2** | 0.0000 | 0.0000 | 77034.0001 | -2.7630 | 0.0057 |
| **layerLoclayer_3** | 0.0000 | 0.0000 | 77034.0001 | 1.1720 | 0.2412 |
| **layerLoclayer_4** | 0.0001 | 0.0000 | 77034.0001 | 8.9364 | 0.0000 |
| **scanTypeSE:layerLoclayer_1** | 0.0000 | 0.0000 | 77034.0001 | 1.7482 | 0.0804 |
| **scanTypeSE:layerLoclayer_2** | 0.0000 | 0.0000 | 77034.0001 | 1.0435 | 0.2967 |
| **scanTypeSE:layerLoclayer_3** | 0.0000 | 0.0000 | 77034.0001 | -0.2576 | 0.7967 |
| **scanTypeSE:layerLoclayer_4** | -0.0001 | 0.0000 | 77034.0001 | -1.6170 | 0.1059 |

##### LRT and PBtest Summary

| Iteration | Tested_Variable | LRT_statistic | LRT_df | LRT_p_value | PB_statistic | PB_p_value |
|---|---|---|---|---|---|---|
| **1** | scanType | 61.7749 | 7.0000 | 0.0000 | 61.7749 | 0.0001 |
| **2** | layerLoc | 13.2940 | 8.0000 | 0.1021 | 13.2940 | 0.1558 |
| **3** | scanType:layerLoc | 5.9310 | 4.0000 | 0.2044 | 5.9310 | 0.2742 |

#### S15: PPU Quality Assessement

| Subject | GE/SE | PPU Quality | Mean BPM | Relative Cardiac Power | Inter Beat Interval Outlier Fraction | Mean Correlation of Cardiac Cycle to Mean Cardiac Cycle | Template Correlation Outlier Fraction |
|---|---|---|---|---|---|---|---|
| **sub-resp02** | GE | BAD | 67.38 | 0.2754 | 0.1659 | 0.8405 | 0.0538 |
| **sub-resp02** | SE | BAD | 71.78 | 0.1681 | 0.3005 | 0.746 | 0.0622 |
| **sub-resp03** | GE | BAD | 96.68 | 0.4715 | 0.1488 | 0.8662 | 0.0706 |
| **sub-resp03** | SE | BAD | 101.07 | 0.2818 | 0.2258 | 0.8462 | 0.0587 |
| **sub-resp04** | GE | GOOD | 49.8 | 0.4304 | 0.0407 | 0.9456 | 0.0392 |
| **sub-resp04** | SE | GOOD | 48.34 | 0.4431 | 0.0485 | 0.9383 | 0.0352 |
| **sub-resp05** | GE | BAD | 62.99 | 0.162 | 0.3598 | 0.7227 | 0.0665 |
| **sub-resp05** | SE | BAD | 175.78 | 0.1392 | 0.4262 | 0.72 | 0.0665 |
| **sub-resp06** | GE | GOOD | 65.92 | 0.5889 | 0 | 0.9875 | 0.0534 |
| **sub-resp06** | SE | GOOD | 67.38 | 0.573 | 0.0033 | 0.9853 | 0.0155 |
| **sub-resp07** | GE | GOOD | 55.66 | 0.6432 | 0.0161 | 0.9731 | 0.0228 |
| **sub-resp07** | SE | GOOD | 52.73 | 0.611 | 0.0094 | 0.9772 | 0.0188 |
| **sub-resp08** | GE | GOOD | 67.38 | 0.5465 | 0.0021 | 0.9826 | 0.0226 |
| **sub-resp08** | SE | GOOD | 62.99 | 0.5562 | 0.0034 | 0.9775 | 0.0101 |
| **sub-resp09** | GE | GOOD | 60.06 | 0.3878 | 0.0153 | 0.9486 | 0.0235 |
| **sub-resp09** | SE | GOOD | 57.13 | 0.3997 | 0.0259 | 0.9386 | 0.0336 |
| **sub-resp10** | GE | GOOD | 68.85 | 0.398 | 0.011 | 0.9553 | 0.0231 |
| **sub-resp10** | SE | GOOD | 60.06 | 0.4388 | 0.0081 | 0.9578 | 0.0127 |
| **sub-resp11** | GE | GOOD | 61.52 | 0.6957 | 0 | 0.9808 | 0.0339 |
| **sub-resp11** | SE | GOOD | 60.06 | 0.7022 | 0.0025 | 0.9798 | 0.005 |
